# Emergent metabolic parasitism driven by organelle sequestration

**DOI:** 10.1101/2025.11.03.686365

**Authors:** Erica Lasek-Nesselquist, Holly V. Moeller, Christopher Paight, Brandon K.B. Seah, Nurislam Shaikhutdinov, Estienne C. Swart, Matthew D. Johnson

## Abstract

Stable acquisitions of metabolism, such as the endosymbiotic incorporation of eukaryotic chloroplasts, are thought to proceed through mechanisms that increase the genetic repertoire of the host and allow for vertical integration of new metabolism. Here we test these predictions using the chloroplast-stealing marine ciliate genus *Mesodinium* by comparing transcriptomes from species that represent a spectrum from full heterotrophy to nearly full phototrophy. In contrast to theory, we find a striking divestment in metabolic autonomy with increased reliance on acquired photosynthesis. Indeed, the highly photosynthetic, red tide-forming *Mesodinium rubrum* appears to have lost the capacity to synthesize amino acids, metabolize fatty acids, and produce peroxisomes. Our results portray a metabolic parasite, masquerading as a free-living ciliate, yet incapable of satisfying most of its basic anabolic needs.

## Introduction

Many eukaryotic lineages rely on acquisitions of metabolic capacity from other species in order to meet their energetic and material needs (*1*). These expansions of metabolism can affect an organism’s contemporary ecology (e.g., by supporting the broadening of the metabolic niche) and may also impact evolutionary trajectories (e.g., by promoting co-evolution or by supporting adaptive radiations) (*2*, *3*). Acquisitions may occur through a variety of mechanisms, including the intracellular integration of endosymbiotic cells or by stealing and retaining functional organelles. However, it remains unclear how the integration of novel metabolic machinery into a new host cell alters the function of the host cell’s genome, especially in terms of metabolic regulation. Major changes in host transcription should reveal instances of dependencies upon foreign metabolic pathways and shed light on early events in the integration of foreign organelles prior to genetic integration.

Kleptoplasty, or the trophic acquisition of functional chloroplasts, is one example of a metabolic acquisition with profound consequences for cellular physiology (*4*). Organisms that transiently obtain the capacity for photosynthesis in this manner are able to expand their metabolic repertoire for energy and carbon acquisition, but must also contend with photooxidative stress from light-harvesting machinery. Nevertheless, dozens of lineages from diverse taxa, including Ciliata, Dinoflagellata, Cryptista, Discoba, Platyhelminthes, Mollusca, and Foraminifera have evolved this capacity (*5–7*), supporting anywhere from 5 to 95% of their cellular carbon demands via acquired chloroplasts (*8*).

The effects of these acquisitions on host genomes remain unclear, in part because strategies for metabolic integration are as diverse as the kleptoplastidic lineages themselves. In some cases, this is supported by genes from either the current kleptoplast lineage or from lost ancestral plastids that have been horizontally integrated into the host nucleus (*7*, *9–11*). In other cases, hosts retain transcriptionally active prey nuclei alongside chloroplasts (*12–14*), and these nuclei continue to regulate plastid function in the new host (*15*, *16*).

Studies of highly integrated kleptoplastidic lineages provide some insights into host gene expression in the context of acquired photosynthesis. Previous work has primarily focused on the extent to which plastid control is integrated into the host genome through the expression of horizontally transferred genes (*7*, *17*, *18*). While some work has sought to contrast expression in heterotrophic and kleptoplastidic lineages (e.g., *19*), studies of how genome function changes during the evolutionary transition towards kleptoplasty are hampered by the genetic diversity of the lineages involved.

### A unique model system

Here, we address this challenge using a comparative approach focused on the *Mesodinium* genus of ciliates. Species within the genus span a gradient of reliance on photosynthesis (*20*, *21*), creating a natural experiment that allows us to compare gene expression as a function of photosynthesis across lineages with a similar genetic background (*3*). We compared: the heterotrophic *M. pulex*, which digests diverse prey immediately upon ingestion, the mixotrophic *M. chamaeleon*, which steals organelles from a variety of cryptophyte species, and the nearly photosynthetic *M. rubrum* which steals organelles from one lineage of cryptophytes. Along this continuum, the more photosynthetic the *Mesodinium* lineage, the fewer prey it ingests, the higher its photosynthetic rate, the greater its complexity of foreign organelle organization, and the more control it retains over kleptoplastids (Fig. 1).

**Fig. 1.**
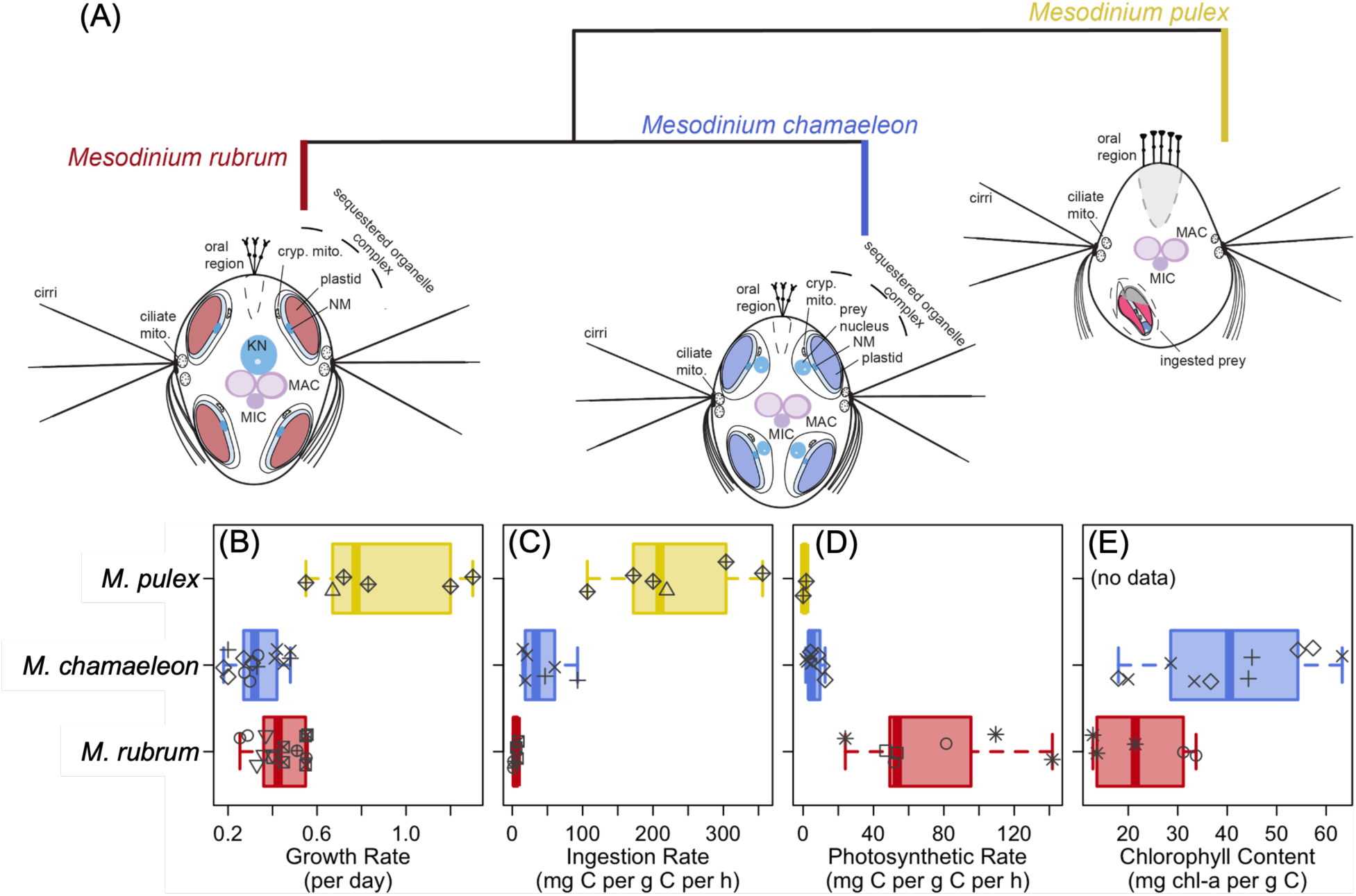
The *Mesodinium* genus spans a gradient of reliance on kleptoplasty. *Mesodinium* (**A**) phylogeny and comparison of (**B**) growth rates, (**C**) prey ingestion rates, (**D**) photosynthetic rates, and (**E**) photosynthetic pigment content across the three major lineages used in this study. Boxplots show ranges of data, and are overlaid with the actual datapoints; shape indicates study from which data are drawn.

In *M. rubrum*, the integration of stolen organelles and their exploitation of prey metabolism is exceptional among organelle stealing organisms: *M. rubrum* can divide its kleptoplastids and grow photosynthetically so rapidly that it forms red water “blooms” visible from space (e.g., *22*– *24*). This remarkable evolutionary feat is made possible by stealing its prey’s nucleus (*12*, *25*), which does not divide in its new host but remains transcriptionally active (*15*, *16*, *26*), and by evolving key innovations in how it organizes foreign organelles. The apparent connection of the “kleptokaryon”, which resides separately from other cryptophyte organelles along with the ciliate nuclei (Fig. 1), with the *Mesodinium* endomembrane system somehow affords the ciliate greater regulatory control over their sequestered organelles. Further, the specialization of *M. rubrum* to a specific subset of cryptophyte species has likely supported their ability to adapt to the products of foreign metabolic pathways and integrate them into their own metabolism.

We hypothesized that increasing reliance on photosynthesis would be accompanied by increased integration of ciliate and cryptophyte metabolic pathways. We found that repeated acquisitions of new nuclei from free-living prey appears to have precipitated a cascading loss of metabolic independence in *M. rubrum* that distinguishes it from its congeners and results in a pattern of gene expression that more closely resembles an endoparasite than a free-living ciliate.

## Results and Discussion

### Loss in protein diversity due to metabolic “parasitism”

To assess evolutionary trends across the *Mesodinium* genus, we assembled seven *Mesodinium* lineage transcriptomes: three from RNA-Seq data that we generated for *M. pulex* strain EPMP20B2, *M. chamaeleon* strain NRMC1802*, and M. rubrum* strain CBJR05, and four from data obtained from NCBI’s Sequence Read Archive (SRA) for *M. chamaeleon* strain NRMC1501 and *M. rubrum* strains MBL-DK2009, Nantes, and CCMP 2563 (table S1). We annotated predicted proteins and assigned pathways via KEGG to compare pathway completeness across lineages. The total number of unique proteins was highest for *M. pulex*, intermediate for *M. chamaeleon*, and lowest for *M. rubrum*, (table S1; fig. S1). While the number of reads sequenced was highly correlated with the number of proteins predicted (Spearman correlation coefficient = 0.82, p-value = 0.03), larger libraries produced more duplication rather than more unique KOs (table S1).

The reduced protein diversity in kleptoplastidic *Mesodinium* lineages was largely a result of reduced numbers of metabolic proteins. While reductive evolution is a characteristic of all life, it is most recognized as a driving force in symbiotic and parasitic organisms, which are thought to be particularly subject to bottlenecks and gene loss due to genetic drift (*27*). Iterative gene loss in intracellular parasites can render them unable to live independently (*28*). When metabolites synthesized by a host are available to a parasite in a stable association, it can lose functionally redundant pathways without a reduction in fitness. As a consequence, many intracellular parasitic lineages, including apicomplexans, trypanosomes, and microsporidia, have been shown to be dependent on their hosts for basic carbon and energy resources, as well as numerous secondary metabolites (e.g., *29*).

In *Mesodinium*, acquired metabolism correlates with similar patterns of loss as seen in apicomplexans and other intracellular parasites. We found that the relative numbers of unique proteins per pathway were similar for genetic information, environmental information processing, and cellular processes across all *Mesodinium* species, but declined with increasing photosynthetic capacity for metabolism (Fig. 2, fig. S2). Our results were robust to transcriptome filtering method (fig. S2), and are likely conservative estimates given that we had more sequencing data and more experimental treatments for our kleptoplastidic lineages than the heterotrophic *M. pulex*, which nevertheless had the highest protein diversity (table S1, fig. S1).

**Fig. 2.**
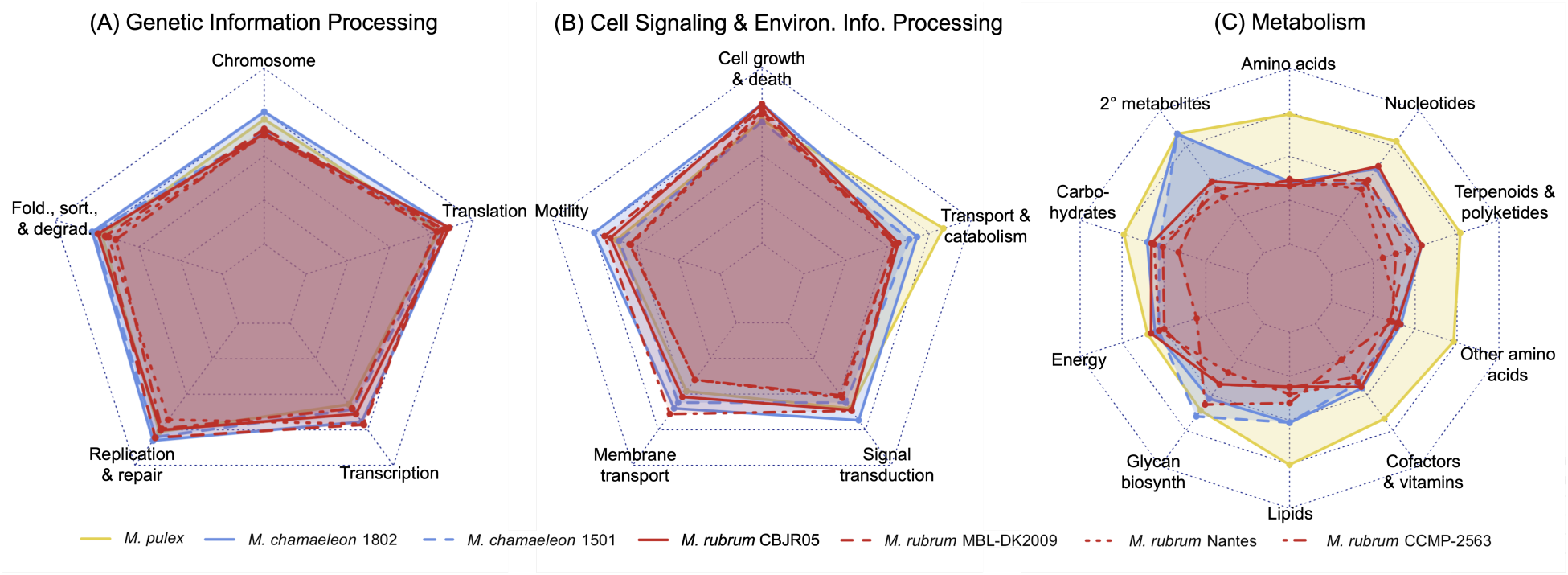
Variation in transcriptome coverage by *Mesodinium* lineage. Axes are scaled by total number of unique proteins in a subcategory across all *Mesodinium* transcriptomes used. For (**A**) genetic information processing and (**B**) cell signalling and environmental information processing, all seven lineages have similar numbers of genes, while for (**C**) metabolic genes, *M. rubrum* lineages are highly depleted.

Patterns of presence/absence across metabolic pathways for *M. rubrum* strains were also consistent with those of the macronuclear genome of *M. rubrum* CBRJ05 (Pearson correlation coefficients ranging from 0.49-0.75, p-values << 0.01). In fact, the *M. rubrum* CBRJ05 genome only contained 14 additional metabolically-related KOs that the *M. rubrum* transcriptomes collectively lacked (two being putative HGTs filtered from transcriptomes). Similarly, read counts for the top 50 most expressed genes in each library did not demonstrate extreme signals (table S2), indicating pathway deficiencies were not an artifact of hyper-abundantly expressed genes biasing transcriptome composition.

We propose that having access to essentially “free” fully synthesized metabolites tapped from sequestered organelle complexes has acted like an evolutionary ratchet, driving the loss of key genes involved in the biosynthesis of certain biomolecules (i.e. certain amino acids, fatty acids, vitamins, and nucleotides). This trend is further reflected in the widespread reduction of redundant and anaplerotic metabolic pathways in *M. rubrum*, which is perhaps most dramatically reflected in their reduced pentose phosphate pathway and amino acid and fatty acid degradation pathways (table S2). For *M. rubrum*, this was true for all subcategories within metabolism except for energy metabolism and glycan biosynthesis and metabolism (Fig. 2C). This reduction in metabolic proteins correlated with reduced connectedness of metabolic pathways to all other KEGG subcategories in *M. chamaeleon* and, especially, *M. rubrum* (Fig. 3, legend in fig. S3).

**Fig. 3.**
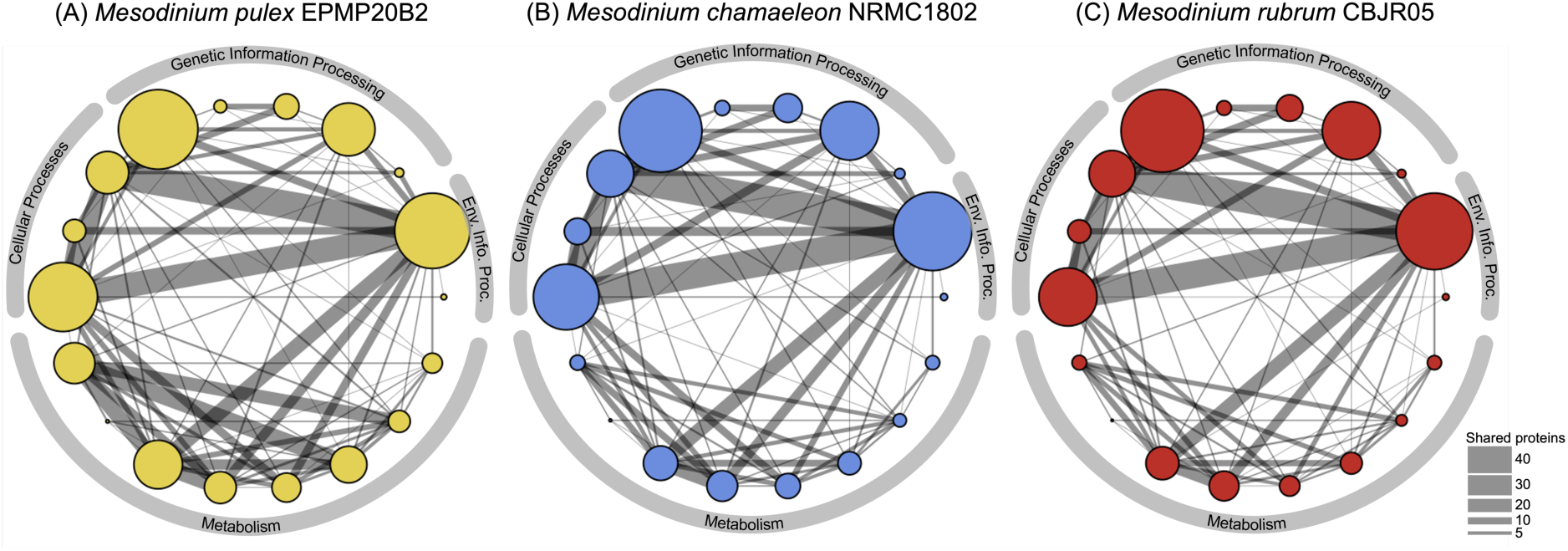
Connectivity of KEGG subcategories within *Mesodinium* lineages. Nodes represent 18 subcategories within four KEGG categories (gray arcs), sized proportionately to number of unique proteins in that subcategory. Network edge line thickness is proportional to number of proteins shared by two subcategories. From *M. pulex* (**A**) to *M. chamaeleon* (**B**) to *M. rubrum* (**C**), reductions in overall pathway completeness also result in reductions in numbers of shared proteins. *M. chamaeleon* data are from culture NRMC-1802, and *M. rubrum* data are from strain CBJR05. See Supplemental Figure 3 for a key to the subcategory identities.

We also observed statistically significant (adjusted p-value < 0.05) reductions in protein diversity within metabolic pathways as lineages increasingly rely on photosynthesis. In *M. chamaeleon*, 26 of 59 metabolic pathways were depleted in protein content in at least one lineage compared to *M. pulex*, and in *M. rubrum*, more than 70% (42 of 59) were depleted in at least one of the four lineages studied (Fig. 4). In contrast, pathways associated with genetic information processing were largely intact (Fig. 4). At the species level, enrichment analyses revealed that 50% of metabolic pathways tested (29 of 58) were collectively depleted in *M. rubrum* in comparison to *M. pulex*, with pathways related to amino acid metabolism showing the greatest reduction (table S2). In fact, of the 18 pathways related to amino acid metabolism tested, only four were not significantly reduced (table S2).

**Fig. 4.**
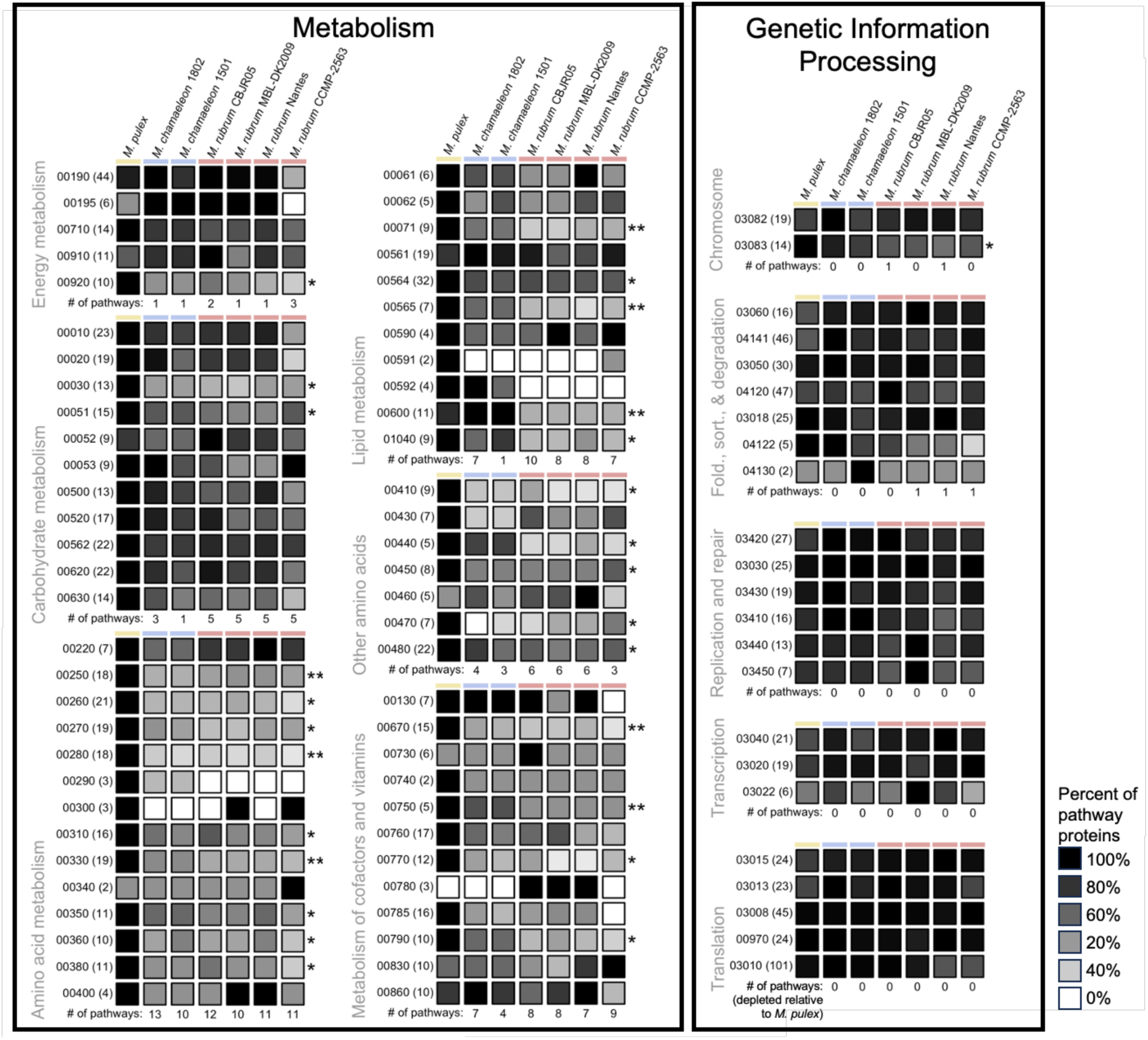
Relative pathway completeness for six Metabolism and five Genetic Information Processing subcategories. Pathways are labeled with KEGG pathway identity and number of unique proteins (KOs) identified in this pathway, and box shading is scaled to the maximum number of proteins in any one *Mesodinium* lineage (black = maximum number of proteins, white = no proteins). Stars indicate significant depletion in *M. rubrum* relative to *M. pulex* (* = p<0.05; ** = p<0.01). Within each subcategory, the number of pathways that are statistically significantly depleted (2.5 standard deviations) relative to *M. pulex* is tallied for each lineage. Note that metabolic pathways are substantially depleted in kleptoplastidic lineages relative to heterotrophic *Mesodinium pulex*, but genetic information processing is largely intact.

Although some variation existed, the specific proteins depleted from each metabolic pathway were largely consistent across *M. rubrum* strains and within *M. chamaeleon* (fig. S4, table S3), supporting the legitimacy of these observations as an evolutionary phenomenon due to their parasitoid-like lifestyle. Further, there was minimal change in the percent completeness of filtered and unfiltered protein datasets (table S1) and all *Mesodinium* species demonstrated approximately the same transcriptome completeness (with the exception of *M. rubrum* Nantes, which had a substantially smaller library size than the other species; table S1). We also noted minimal rRNA contributions to our libraries (table S1), except for *M. rubrum* DK2009, which still showed an equivalent number of unique KOs as compared to *M. chamaeleon* NRMC1501. Thus, rRNA biases likely do not influence the reduction in the number of unique KOs observed for kleptoplastidic *Mesodinium* species. Collectively, these results suggest that neither biased filtering nor over-filtering is responsible for the differences observed between heterotrophic and kleptoplastidic *Mesodinium* species.

### Loss of anabolic autonomy and compensatory expression

Pathways related to lipid metabolism, particularly fatty acid metabolism, were also collectively depleted in *M. rubrum* compared to *M. pulex* (Fig. 2, Fig. 4, table S2). Both of the organelle stealing ciliates, *M. chamaeleon* and *M. rubrum,* appear to have lost the capacity to make isoleucine, leucine, ornithine, tyrosine, and valine compared to their heterotrophic relative *M. pulex* (table S2). Some parallels can be drawn to apicomplexans and the evolution of the parasitophorous vacuole (PV). In *T. gondii*, the recruitment of host proteins to the PV, helps to support numerous metabolic auxotrophies by digesting host metabolites, including the amino acids arginine, tyrosine, and tryptophan (*29*).

When considering fatty acid biosynthesis, degradation and biosynthesis of amino acids, and synthesis of secondary metabolites, the greatest loss is found in *M. rubrum*-consistent with loss being driven by an increasing dependency upon acquired metabolism. This assumption is strongly supported by not only the conspicuous absence of genes comprising these pathways in *M. rubrum* but also their presence within the kleptokaryon (fig. S5, table S3), and the high levels of their metabolic products synthesized in the ciliate (*16*). This metabolic chimerism in mixotrophic *Mesodinium* species helps to explain their obligate requirements for kleptoplasty, and perhaps even their specific requirements for certain cryptophyte species (e.g., *30*, *31*). More broadly, such obligate requirements for photosynthesis in other kleptoplastidic species may also be driven by similar losses, albeit less extreme, in their ability to synthesize key metabolites.

### Loss of peroxisome function

Peroxisome related proteins demonstrated the most conspicuous absence in *M. rubrum*, with the peroxisome being one of the most significantly under-represented categories for this group (table S2). To further contextualize the observed reduction in peroxisome pathways, we compared our *Mesodinium* species to other alveolates (*32*). The loss of peroxisome and peroxisome-related proteins in *M. rubrum* was as extreme or more extreme than most apicomplexan parasites (Fig. 5, fig. S6). The apicomplexans *Plasmodium* spp., *Gregarina niphandrodes* and *Cryptosporidium parvum* have lost most of their peroxisome biosynthetic genes (i.e., peroxins) and much of the classic metabolic genes, such as those involved in beta oxidation, associated with this organelle (*32*, *33*). This reductive evolution in certain apicomplexans is thought to be driven by the reliance of these parasites on their host lipid metabolism pathway (*34*). The significant reduction of amino acid and fatty acid degradation pathways in *M. rubrum* is also likely correlated with the extreme loss of peroxisome-specific proteins, and suggests the absence of a functional peroxisome in *M. rubrum*. The extreme protein reduction associated with the peroxisome in *M. rubrum* was not observed in other cellular processes, such as those related to endocytosis, the lysosome, and the phagosome (tables S1-S2).

**Fig. 5.**
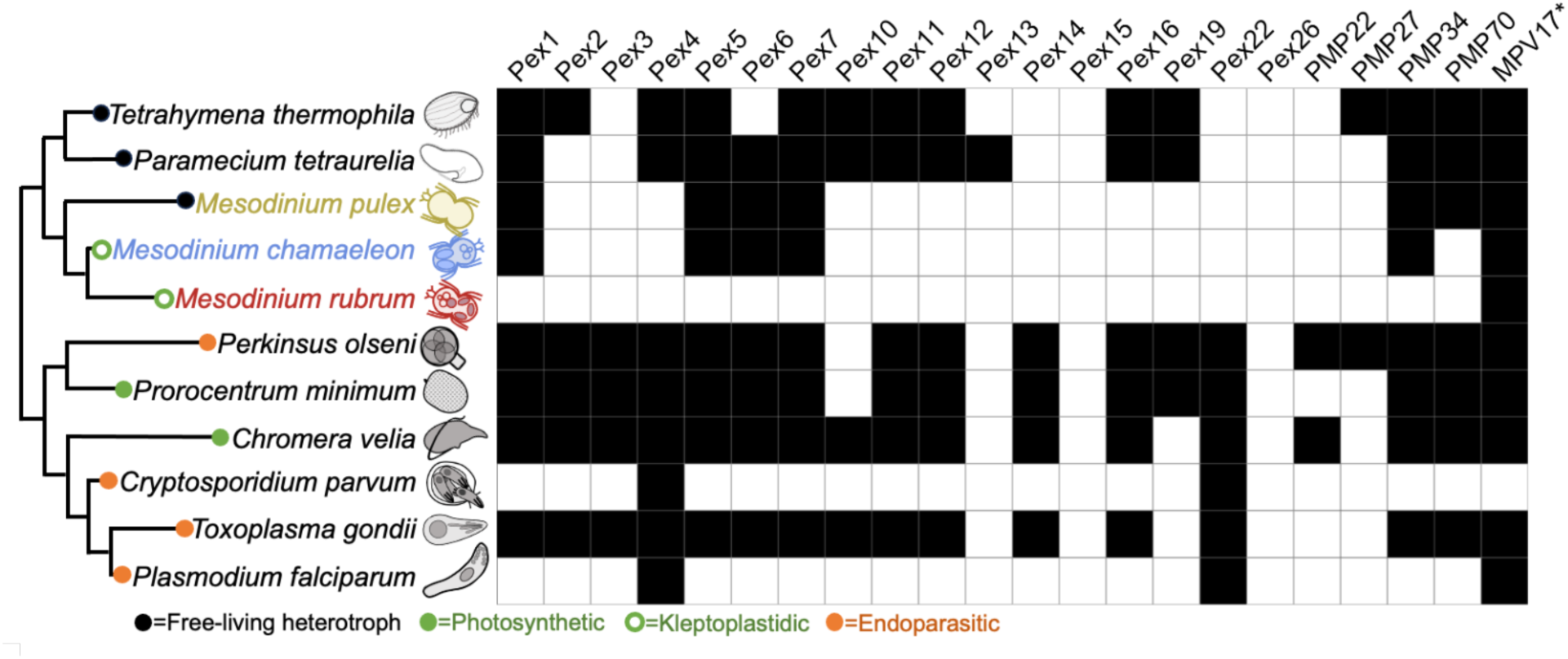
Comparison of peroxisome proteins in *Mesodinium* ciliates and other alveolates. Filled boxes indicate that a protein is present in a species. Proteins classified as peroxisomal are according to Ludewig-Klingner et al. (*32*), and data are from this study or (*32*). The metabolic lifestyle of each lineage is indicated with a colored circle, and evolutionary relationships are shown using an unrooted dendrogram. *M. chamaeleon* data are for culture NRMC-1802, and *M. rubrum* data are for strain CBJR05. *MPV17 is now thought to be a mitochondrial inner membrane protein (*37*).

*Mesodinium rubrum* lacked proteins related to the peroxisome membrane and its organization as well as proteins involved in protein and metabolite importation and translocation. Ciliates from the order Oligohymenophera, as well as *M. pulex* and *M. chamaeleon*, had a more complete repertoire of peroxisome proteins; the latter two were the only *Mesodinium* species with evidence of peroxisomal targeting signals (the C-terminal tripeptide, PTS1 and the N-terminal nonapeptide, PTS2, which mediate protein import), indicating that the loss observed in *M. rubrum* was not a general ciliate or *Mesodinium* genus phenomenon (Figure 5). Additionally, it is possible that the few peroxisome-related proteins retained by *M. rubrum* are actually found in other pathways and/or cellular environments (*35–37*). This loss in metabolic potential fits with the observed loss of fatty-acid degradation (e.g., beta-oxidation) genes in *M. rubrum*, and suggests that most of their fatty acid and lipid biosynthesis occurs in sequestered organelle complexes. This pattern also supports the complementary overexpression of genes in lipid and fatty acid biosynthesis and degradation pathways found in the kleptokaryon of *M. rubrum* (*16*). In cryptophytes, peroxisomes have been shown to be important for metabolism of carbohydrates, ether phospholipids, nucleotides, vitamin K, reactive oxygen species, amino acids, and amines, while beta oxidation of fatty acids occurs in the mitochondrion (*38*).

### Evolutionary implications of metabolic parasitism

Conceptually, parasites and parasitoids are generally considered to require a host as an encompassing habitat. Here we show that in the case of *Mesodinium* ciliates, and perhaps other kleptoplastidic organisms, the parasite concept can be expanded to include “inverted” parasitism, whereby the host-parasite relationship is inverted or turned inside out. Organelle-stealing *Mesodinium* ciliates function as “inverted parasitoids” that kill their prey but continue to exploit their organelles and metabolism, powered by continued gene expression from a stolen nucleus. Selective pressures on parasites, which include metabolic and cellular reduction while maintaining sufficient cell division, has led to convergent patterns among disparate parasitic lineages for gene loss and genome reduction and or compaction (*28*, *39*). Such reductions include both the nuclear and organelle genomes and sometimes results in organelle or organelle genome loss (*40*, *41*). Metabolic functional loss is a common theme in obligate intracellular parasites, such as the loss of de novo nucleotide synthesis in certain apicomplexans (*42*) or the nearly complete loss of genes involved in glycolysis within microsporidians (*43*). We have shown that similar selection mechanisms have both streamlined central metabolic pathways, and resulted in loss of essential biosynthetic capabilities in *M. rubrum*, much like obligate intracellular parasites.

This concept is a new perspective for kleptoplastidic organisms, but one that is more apt than the traditional parallels drawn to endosymbiosis. Rather than establishing a mutualism, organelle thieves kill their prey and enslave their organelles, thereby removing any coevolution with their prey beyond those evolved from predation dynamics. In contrast to parasites, *M. rubrum* ciliates likely do not experience pronounced population bottlenecks, but rather, along with *T*. *amphioxeia*, are known to form massive red-water blooms that can stretch for hundreds of kilometers (*24*). Relative to other ciliates, *Mesodinium* occurs at high population densities and has high potential for interaction due to its behaviors of forming thin layers, “swarms”, and blooms (*44*, *45*). Further, unlike viruses, predators, or parasites, *M. rubrum* does not appear subject to so-called “Red Queen Hypothesis” dynamics, where host-parasite or predator-prey relationships are shaped by a coevolutionary “arms race” (*46*, *47*). Rather, *M. rubrum* is better described by “Black Queen Hypothesis” dynamics, which describes a theory of reductive evolution driven by community dependencies upon “leaky helpers” (*48*). In such scenarios, community members or symbionts may lose biosynthetic genes if they can exploit community or host metabolites. In this instance, sequestered organelle complexes in the ciliate function as leaky “zombie” helpers that sustain their host and reduce its need to be anabolically self-sufficient. We suggest that the apparent gene loss we have shown in *M. rubrum* is driven by reductive evolution due to metabolic dependencies, akin to genetic drift seen in many parasites. The inverted parasitoid paradigm we propose further challenges the mutually beneficial nature of nascent endosymbioses (*49*, *50*), by transcending organismal coevolutionary relationships through organelle “symbiosis”. This unique ciliate clade provides a mechanistically novel model system for investigating early events of a plastid acquisition by a non-photosynthetic lineage, that appears to be guided by new rules.

## Acknowledgments

We thank collaborators at the Department of Energy Joint Genome Institute for providing technical support for RNA sequencing. ELN thanks the Bioinformatics Core at the Wadsworth Center, NYSDOH. Research was sponsored by the United States Army Research Office and was accomplished under Cooperative Agreement Number W911NF-19-2-0026 (to HVM) for the Institute for Collaborative Biotechnologies.

## Funding

Department of Energy Joint Genome Institute Community Sequencing Project 503035 (ELN, HVM, ECS, MDJ)

National Science Foundation grant 2344641 (HVM)

National Science Foundation grant 2344640 (MDJ)

United States Army Research Office W911NF-19-2-0026 (HVM)

## Author contributions

Each author’s contribution(s) to the paper should be listed [we encourage you to follow the <u>CRediT</u> model]. Each CRediT role should have its own line, and there should not be any punctuation in the initials.

Examples:

Conceptualization: ELN, HVM, MDJ

Methodology: ELN, HVM, BKBS, MDJ

Investigation: ELN, HVM, CP, BKBS, MDJ

Visualization: ELN, HVM, MDJ

Funding acquisition: HVM, MDJ

Project administration: HVM, MDJ

Supervision: ELN, HVM, MDJ

Writing – original draft: ELN, HVM, MDJ

Writing – review & editing: ELN, HVM, CP, BKBS, NS, ECS, MDJ

## Data and materials availability

RNASeq data for *M. chamaeleon* NRMC1501, *M. rubrum* nantes, *M. rubrum* DK2009, and *M. rubrum* CCMP2563 (including read data for *Geminigera cryophila* prey) are available in NCBI under the BioProjects: PRJNA560220, PRJEB53438, PRJNA880267, and PRJNA560206, respectively. RNASeq data generated in this study for *M. chamaeleon* NRMC1802, *M. rubrum* CBJR05, *Storeatula major*, and *Teleaulax amphioxeia* GCE-P01 are available through the JGI Genome Portal https://genome.jgi.doe.gov/portal/ under the proposal ID 503035. RNASeq data for *M. pulex* are deposited in NCBI under the BioProject: PRJNA1225163 (reviewer link: https://dataview.ncbi.nlm.nih.gov/object/PRJNA1225163?reviewer=pgkr88s5527erqsk32s0hmhq91). Transcriptome assemblies, predicted coding sequences, protein annotations, and other associated data are available in Dryad: https://doi.org/10.5061/dryad.1jwstqk4x (reviewer link: http://datadryad.org/stash/share/s3Uh8hHzyr0bUMOftrqKg2miXVxD6l4VVaQp0YtWI5E). Code and associated data files for statistical analyses and figure generation are available at https://github.com/hollymoeller/MesodiniumComparativeTranscriptomes.

## Supplementary Materials

### Materials and Methods

#### Generation of transcriptomic data

For this study, we generated new transcriptomic data for three *Mesodinium* species. Heterotrophic *M. pulex* (EPMP20B2) cells were grown in sterile filtered seawater (FSW) with *Storeatula major* as prey, added weekly with fresh media at a 10:1 prey:predator ratio. Cells were cultured in 250 mL tissue culture flasks and harvested for RNA extraction by gently concentrating the ciliates atop of 8.0 μm Transwell filters in 6 well plates (*51*, *52*). About 100 mL of cells were concentrated over each filter with prey and bacteria allowed to pass through, and the *M. pulex* were then washed with 100 mL of FSW before collecting them on a 1 μm polycarbonate filter, freezing with liquid nitrogen, and storing at −80°C until extraction.

Mixotrophic *M. chamaeleon* (NRMC-1802) cells were also grown in FSW with *S. major* as prey, and maintained at a light level of 20 μmol quanta m^-2^ s^-1^ with a 12 hr light: 12 hr dark diel cycle. To maximize the diversity of expressed genes, we collected cells under eight experimental treatments, mostly related to feeding history. In particular, we conducted a 4-week-long experiment in which *M. chamaeleon* cells were first starved for two weeks (“starved”), and then either starved for a further two weeks (“extended starvation”) or pulse-fed at a 10:1 *S. major* to *M. chamaeleon* ratio and sampled 24, 48, 96, and 168 hours post feeding. We also cultivated well-fed *M. chamaeleon* cells under high light (100 μmol quanta m^-2^ s^-1^) and in FSW amended with ammonium.

Highly photosynthetic *M. rubrum* (CBJR05) was cultivated in f/2-Si media (*53*) with *Teleaulax amphioxeia* GCEP01 (*51*) as prey. As with *M. chamaeleon*, we maintained cultures at 20 μmol quanta m^-2^ s^-1^ and used multiple (seven) experimental treatments to maximize the diversity of expressed genes. We collected transcriptomic samples from *M. rubrum* cells that were well-fed *T. amphioxeia* (“control”) and collected either at the midpoint of their day (“noon”) or night (“midnight”) diel cycle. We contrasted these treatments with a starved culture (which was not fed for two weeks prior to collection), and *M. rubrum* grown in different nitrogen environments (without any form of N added to the f/2-Si media, with only ammonium, and with both ammonium and nitrate). Both *M. chamaeleon* and *M. rubrum* samples were collected in the same way as *M. pulex* (washed free of prey using a 8.0 μm transwell filter, followed by collection on a Millipore Omnipore JMWP 5 μm polycarbonate filter and immediate freezing in liquid nitrogen).

So that we could identify and remove contaminating prey RNA sequences, we also collected transcriptomes for *S. major* and *T. amphioxeia*. For both cryptophytes, samples were collected from cells maintained under either high light (100 μmol quanta m^-2^ s^-1^) or low light (10 μmol quanta m^-2^ s^-1^) either at mid-day (6 hrs in to the 12-hr light cycle) or mid-night (6 hrs in to the 12-hr dark cycle).

We extracted RNA from flash-frozen samples using a Qiagen RNeasy Plant kit. For *M. rubrum* CBJR05, RNA was further purified using a lithium chloride precipitation (*54*). Samples were sequenced at the UC Davis Genome Center (*M. pulex* EPMP20B2) or by the DOE Joint Genome Institute Sequencing Center (*M. chamaeleon* NRMC-1802, *M. rubrum* CBJR05, and both cryptophytes).

#### *Mesodinium* transcriptome assembly

Transcriptome assemblies from Illumina data were generated for three *Mesodinium* species *Mesodinium pulex*, *Mesodinium chamaeleon*, and *Mesodinium rubrum*, which included one strain of *M. pulex*, two different cultures of *M. chamaeleon*, and four strains of *M. rubrum* for a total of 7 transcriptomes (table S1). The number of libraries/treatments varied, with *M. chamaeleon* NRMC1802 and *M. rubrum* CBJR05 subject to the most robust sampling - having the greatest number of treatments and replicates per treatment (table S1). Illumina sequencing was performed by this study or obtained from the SRA database (table S1). Prey transcriptomes were obtained from JGI or assembled with Megahit v.1.2.9 when data were available.

All *Mesodinium* libraries were processed and assembled by an in-house Nextflow v.2 workflow. TrimGalore v.0.6.7 (https://github.com/FelixKrueger/TrimGalore) removed adapters from raw reads, excluded reads < 75 bp in length, and trimmed the ends of reads with a PHRED quality score < 20. Reads classified as bacterial by Kraken 2 v.2.1.3 (*55*) with the standard-8 (https://benlangmead.github.io/aws-indexes/k2) database were removed. Prey contamination was removed by mapping bacteria-filtered reads to the prey transcriptome with BBmap v.39.01 (https://sourceforge.net/projects/bbmap/) using the following parameters: minratio=0.9, maxindel=3, bwr=0.16, bw=12, minhits=2, qtrim=r, trimq=10, kfilter=25, maxsites=1 and k=14 (summary statistics: table S1).

After bacteria and prey removal, the remaining reads were assembled with Megahit v.1.2.9 (*56*) using the “meta-large” parameter and clustered at 97% identity by cdhit-est v.4.8.1 (*57*, *58*) to reduce redundancy. Transcripts classified as bacterial by Kraken 2 (*55*) using the standard-8 database were removed. Prey contamination at the transcript level was removed by Blastn v.2.14.0+ (*59*) searches against a cryptophyte database containing transcripts from all prey assemblies generated during this study as well as all cryptophyte genomes available from NCBI’s GenBank as of 2022. *Mesodinium* transcripts with >= 75% identity and >= 75% query coverage to a hit in the cryptophyte database were excluded. Transcripts from each library/treatment were concatenated into a single file for each *Mesodinium* sample and redundancy was reduced by performing another round of clustering with cdhit-est at 97% identity (table S1).

Transdecoder v.5.7.1 (Haas, BJ. https://github.com/TransDecoder/TransDecoder) identified putative coding regions using both the TransDecoder.LongOrfs and TransDecoder.Predict functions and the *Mesodinium* genetic code (#29 in NCBI’s “The Genetic Codes”). Protein predictions from cryptophyte transcriptomes were conducted in the same manner with the standard genetic code. Final protein predictions were clustered at 95% identity by CD-HIT. Proteins were annotated by the KEGG server (https://www.kegg.jp/) using the GhostKOALA option (*60*).

#### Contamination removal at the protein level

Bacteria and prey contamination were removed at the protein level (table S1) by Diamond v.2.1.8.162 (*61*) blastp searches against the NCBI non-redundant (nr) database (downloaded 2024-04-24) and a custom database containing proteomes from representatives of major eukaryotic lineages (archaeplastida, metazoa, fungi, dinoflagellates, stramenopiles, rhizarians, apicomplexans, cryptophytes, euglenids, and haptophytes), all available ciliate proteomes from GenBank as of 2022 (*Blepharisma stoltei*, *Halteria grandinella*, *Ichthyophthirius multifiliis*, *Moneuplotes crassus*, *Oxytricha trifallax*, *Paramecium octaurelia*, *Paramecium pentaurelia*, *Paramecium primaurelia*, *Paramecium sonneborni*, *Paramecium tetraurelia*, *Paramecium tetraurelia* strain d4-2, *Pseudocohnilembus persalinus*, *Stentor coeruleus*, *Stylonychia lemnae*, *Tetrahymena thermophila* SB210), other alveolates (alveolates are defined here as ciliates, apicomplexans, copodellids, and colponemids but not dinoflagellates as many have a photosynthetic history with nuclear contributions from red algal lineages), Archaea, and Bacteria predominantly from the order *Alphaproteobacteria* but also including members of *Gammaproteobacteria* and *Bacteriodota*, which were initially identified as common contaminants in *Mesodinium* transcriptomes by GhostKOALA results.

Proteins assigned KO numbers by GhostKOALA were searched against the nr and custom databases with the default parameters of Diamond and the number of hits returned for each query set as 25 and 50, respectively (table S4). Comparing results between the nr and custom databases revealed that the high number of bacterial sequences in the nr database sometimes prevented the identification of equally good hits to eukaryotes, despite increasing the number of hits returned. This was rectified by the more equal taxonomic sampling present in the custom database. Conversely, the sometimes inadequate bacterial sampling in the custom database resulted in ambiguous taxonomic sequence assignments, which were rectified by the presence of strong hits to bacteria (>=90% identity and >=90% query coverage) in the nr database. Finally, the number of cryptophyte sequences in the custom database greatly exceeded that of the nr database, which helped flag cryptophyte contamination not identified by the nr database.

Developing and testing contaminant filtering parameters was critical to downstream analyses. Ciliates are taxonomically undersampled, with only 149 assemblies listed in NCBI as of 2025-02-04. The majority of these are representatives of the ciliate classes Oligohymenophorea and Spirotrichea, which are separated from the Litostomatea to which *Mesodinium* belongs, by ∼1000 million years (*62*). In fact, the median percent identity (pid) observed between *Mesodinium* sequences and alveolate hits was only 40%. Available litostome genome assemblies predominantly derive from rumen ciliates and protein prediction and annotation suggested that these assemblies contained bacterial contamination. Furthermore, *Mesodinium* has been noted as having an accelerated rate of evolution (*63*), compounding the difficulty of identifying legitimate ciliate sequences in our mixed-species transcriptomes. Cryptophytes are secondary endosymbionts with red alga-derived plastids and remnant algal nuclear genomes (nucleomorphs). The mosaic composition of the cryptophyte can distort phylogenetic signals and further complicate contamination removal. Thus several factors were considered to ensure that filtering was stringent enough to discard most false positives without also removing potential *Mesodinium* sequences that lacked clear homology to those of other ciliates.

The GC content of each coding sequence was calculated by the geecee function from EMBOSS v.6.6.0.0 (*64*). Appropriate filtering cutoffs based on GC content were established by comparing the GC distributions of prey coding sequences to those of sequences in *Mesodinium* transcriptomes that returned best blast hits to alveolates in both nr and custom databases. GC summary statistics were calculated in base R and distributions were plotted with the ggplot2 package. The median GC value of 0.6 for cryptophyte coding sequences was set as a reasonable cutoff for identifying contaminants given that despite the strikingly different median GC value of *Mesodinium* sequences (0.39), there was substantial overlap in prey and host GC distributions (fig. S7).

Alien indices were calculated for any sequence that had a non-alveolate best blast hit in either the nr or custom database with the following formula: log10(e-value of best alveolate hit + 1e-200) −log10(e-value of best blast hit + 1e-200) (*65*). The alien index was set to “na” if the custom database failed to return a hit to an alveolate or all hits returned were to a single taxonomic group for the nr database. Although typically employed to identify horizontal gene transfers (*65*), the alien index here provided a measure of equivalency between non-alveolate best blast hits and alveolate hits for each query sequence, with an alien index of zero indicating an equally good alveolate hit and a larger alien index supporting the non-alveolate classification of the sequence.

Appropriate filtering cutoffs based on alien indices were established by analyzing *Mesodinium* sequences that had a best blast hit to an alveolate in one database but not the other. Sequences identified as true contaminants by additional blast searches and phylogenetic placement had alien indices >= 25 (these were removed from final analyses). Alien index cutoffs >= 10 and >= 50 were tested but the first filtered too aggressively while the second filtered too leniently; the latter being five orders of magnitude above the initial cutoff employed for detecting HGT (*65*). Others have defined an alien index >= 30 as strong evidence of HGT (*66*, *67*). This suggests that an alien index of >= 25 will be highly specific in identifying contamination but not sensitive enough to remove all non-ciliate sequences.

#### Filtering strategies

Any proteins that lacked a hit to the nr database or proteins that returned a top hit with an e-value >= 1e-05 were excluded from further analysis. Any protein with a non-alveolate best blast hit that shared a KO assignment with a protein that had an alveolate best blast hit in either database were not evaluated. Any sequence with a best blast hit to an alveolate in either the nr or custom database were always retained, regardless of the filtering strategy. The remaining sequences were subject to three filtering strategies (<u>doi.org/10.5061/dryad.1jwstqk4x</u>), with the first being the least permissive.

Strategy 1 removed any sequence with an alien index >= 25 in either database and sequences that failed to return any hit to an alveolate sequence as contamination. This essentially retained sequences with strong alveolate signals or those with homology to alveolate sequences present in the nr or custom databases.

Strategy 2 removed any sequences with an alien index >= 25 in either database but further evaluated sequences without any alveolate hits. These sequences were considered contamination if they returned a best blast hit to bacteria, archaea, or algae (defined here as stramenopiles, dinoflagellates, cryptophytes, the archaeplastida, rhizaria, euglenids, and viridiplantae) with a pid and percent query coverage (qcov) >= 70% or with a GC content >= 0.60. Sequences were also removed if the nr and custom databases concordantly returned bacteria or cryptophyte best blast hits. This strategy acknowledges that *Mesodinium* spp. likely harbor proteins lacking homologs in other alveolates and that the similarity between *Mesodinium* and alveolate homologs is often low, increasing the likelihood that *Mesodinium* proteins will return spurious hits to other species. However, a pid >= 70% to a bacterial or algal sequence is well above the median shared homology between *Mesodinium* and alveolate proteins. The requirement that *Mesodinium* proteins also share >=70% qcov with bacterial or algal proteins before removal likely results in conservative filtering. Concordance between nr and custom databases was considered as reasonable support for the identification of bacterial and cryptophyte contamination but insufficient upon manual inspection of the results and further phylogenetic placement of select sequences. Nevertheless, the patterns of protein presence/absence across pathways remained consistent with Strategy 1 and Strategy 3 (fig. S2).

Similar to Strategy 2, Strategy 3 removed any sequences with an alien index >= 25 but further evaluated sequences without alveolate hits. These sequences were removed if they returned a best blast hit to an algal sequence with pid and qcov >= 70%, a best blast hit to a cryptophyte in both databases, or if the sequence had a GC content >= 0.60 with a top hit to a bacterial or algal sequence. Additional prokaryotic contamination was filtered when sequences returned best blast hits to bacteria/archaea in either database with a pid >= 50% or when 100% of the blast results or one less than the total number of blast results returned for both databases were to bacteria/archaea. Sequences that returned a best blast hit to bacteria in the nr database but a non-algal hit in the custom database were also removed if the pid of the bacterial hit was >= 25% that of the hit from the custom database. These cutoffs were selected to balance removing bacterial and algal contamination while retaining putative *Mesodinium* proteins with no alveolate affiliation.

#### Transcriptome completeness

Busco v5.8.0 (*68*) evaluated transcriptome completeness at the protein level for annotated proteins prior to and after the contamination removal using the “alveolate_odb10” database.

Transcriptomes were also compared to the macronuclear genome of *M. rubrum* CBJR05 (table S2). The *M. rubrum* genome assembly and analysis will be described in a separate study. Briefly: To purify macronuclei (MACs), cells were lysed with Galbraith’s solution (Galbraith et al., 1983) to release nuclei, which were fixed with 0.2% (w/v) formaldehyde overnight. Fixed nuclei were resuspended in 3% BSA/TBSTEM, and labeled with rabbit anti-H3K9ac (Merck 06-942, 1:100 dilution) primary antibody, and goat anti-rabbit Alexa 488 conjugate (Abcam ab150077, 1:200 dilution) secondary antibody. Immunofluorescence-labeled nuclei were stained with DAPI, resuspended in Galbraith’s buffer, and flow-sorted on a BD FACSMelody instrument gated on DAPI and Alexa 488 fluorescence (flow-sorting protocol adapted from *69*). Sorted nuclei populations corresponding to MACs were verified by microscopy. Extracted MAC gDNA was sequenced by PacBio HiFi. Reads were assembled with Flye v2.9.2 (*70*). Contigs >60% GC were excluded, as they comprised bacterial contaminants.

#### rRNA quantification and read distribution

The rRNA content of each library was evaluated by mapping reads to 18s rDNA sequences with Salmon v.1.10.3 (*71*) (<u>doi.org/10.5061/dryad.1jwstqk4x</u>). Representative genes were obtained from the nucleotide database of NCBI for all species and specifically for the *M. rubrum* strains CBJR05, CCMP 2563, and DK2009 (accession numbers: JN084213.1, DQ411865.1, JN412738.1, KX783613.1, and MG018339.1). Similarly, Salmon quantified the expression of all *Mesodinium* spp. coding sequences, and the top 50 most expressed genes from each treatment/replicate were used to evaluate read distribution within a transcriptome (table S5). The evenness of read counts across transcripts was summarized by calculating Pielou’s Evenness Index. Briefly the read counts associated with each transcript were divided by the total number of reads in the sample. These proportions (p) were multiplied by their natural logarithms ln(p) and all products were summed to generate the Shannon Diversity Index (H). The Shannon Diversity Index was divided by the natural logarithm of the sample size (ln(S)), resulting in Pielous’s Evenness Index. An index closer to one indicates a more even community representation or in this case a more even distribution of reads among transcripts.

#### Kleptokaryon transcriptome assembly

To evaluate the pathway completeness of the kleptokaryon in relation to its host, assemblies were generated by Megahit for *M. chamaeleon* NRMC1802 and *M. rubrum* CBJR05 with reads filtered for bacterial but not cryptophyte contamination. Transcripts were aggregated at the 97% identity level by cd-hit and subject to another round of bacterial contamination removal as performed previously. Similarly, Transdecoder was employed to predict proteins, but with the standard genetic code, and proteins were annotated by GhostKOALA. Annotated proteins were queried against nr and custom databases by Diamond and only queries returning tophits to cryptophytes were retained.

#### Rarefaction analyses

To generate rarefaction curves, transcripts quantified by Salmon were associated with KO numbers. A transcript was required to have a read count >= 10 to be counted as an occurrence of a KO in a library. The number of occurrences of a KO were aggregated by treatment for each transcriptome (table S6). The resulting tables of KO counts were imported into R and rarefaction curves were generated by the rarecurve function of the *vegan* package (*72*).

#### Statistical analyses

We first assessed the relative numbers of genes across KEGG categories by tallying all unique KO numbers per transcriptome. We normalized the number of genes in a lineage to the total number of unique genes found across all 7 *Mesodinium* transcriptomes to scale the axes in Fig. 2.

Having identified Genetic Information Processing (GIP) as the KEGG category with the greatest completeness and evenness across the *Mesodinium* lineages, we used expression in this category to set a baseline of comparison for other categories and pathways. We first calculated the mean number of genes across all 7 *Mesodinium* transcriptomes for each pathway within the GIP category. Next, we normalized the pathway gene counts of each of the 7 lineages to these means. Thus, for each pathway, we then had a distribution of relativized completeness scores centered around 1. We next calculated the mean and standard deviation of relativized completeness across GIP pathways for each *Mesodinium* lineage. Note that standard error (a more conservative metric than standard deviation) overlaps 1 for GIP for all lineages (fig. S2), indicating the utility of GIP pathways as a common reference point.

We interpreted the standard deviation of GIP pathway completeness (SD_GIP_) as a metric of variance in each *Mesodinium* lineage. Because we expected high evenness (and therefore relativized pathway scores of 1) for GIP pathways, deviations are an indication of sequencing and analysis noise. Therefore, we used SD_GIP_ as a tool to score over-or under-representation of genes in pathways in other KEGG categories. To do so, we first relativized each lineage’s gene count within each pathway by the mean gene count for that pathway across all 7 lineages (as with GIP). We then subtracted 1 from these relativized counts, such that over-representation had positive scores and under-representation had negative scores, and divided by SD_GIP_. Our data were sequenced on different platforms by different laboratories, so we chose to use lineage-specific SD_GIP_ metrics for these calculations, but note that these values are similar and more conservative than genus-level SD_GIP_. The result is a metric that reports, in number of standard deviations, the amount by which a particular *Mesodinium* lineage’s pathway has relatively more or fewer expressed genes than the genus average.

#### Pathway enrichment calculations

Pathway enrichment scores were calculated by dividing the proportion of unique proteins (KO numbers) for each *Mesodinium* spp. per pathway by the proportion of total unique proteins assigned to a species. The proportion of unique proteins per pathway represents the number of unique proteins assigned to that pathway per species/strain divided by the total number of unique pathway proteins across all *Mesodinium* species. The proportion of total unique proteins per species represents the total number of unique KO numbers assigned to a species divided by the total number of unique KO numbers identified across the entire dataset (all *Mesodinium* spp.). Mean pathway fold enrichment scores for *M. rubrum* samples were compared to those of *M. pulex* with a one-sample Student’s t-test, where *M. pulex* represented the population. FDR-adjusted p-values were calculated in R v.4.4.1 with the p.adjust function (table S2).

#### Network analyses

Network analyses depicting the degree of connectedness among KEGG subcategories for *M. pulex*, *M. chamaeleon NRMC1802*, and *M. rubrum* CBJR05 were visualized with the igraph package (*73*) for R. KEGG subcategories (nodes) were weighted by the number of unique KO numbers present and were connected to other subcategories by shared KO numbers (edges).

#### Pathway correlation analysis

Presence/absence data for KOs related to metabolism across *Mesodinium* transcriptomes and the *M. rubrum* CBRJ05 macronuclear genome were represented in a binary table (table S2) and imported into R v.4.4.1. Pearson correlation coefficients and statistical significance were calculated and exported as an Excel document by the cormatrix_excel function in the rempsyc package (*74*).

#### Peroxisomal targeting sequences

Single-line sequences of peroxisome-associated proteins were analyzed for peroxisomal targeting signals type 1 (PTS1) and 2 (PTS2) with regex grep commands. PTS1 tripeptide signals were identified by searching for the consensus sequence [S,A,C]-[K,R,H]-[L,M] (*75*) at the C-terminal end of each protein. The much rarer N-terminal PTS2 nonapeptide was searched for in the first 50 amino acids of each protein with the traditional consensus sequence [RK]-[LVI]-X5-[HQ]-[LA] (used by PSORT II) and the more specific consensus sequence [R/K]-[L/V/I/Q]-X2-[L/V/I/H/Q]-[L/S/G/A/K]-X-[H/Q]-[L/A/F] proposed by Petriv et al. (*76*).

---

**Fig. S1.**
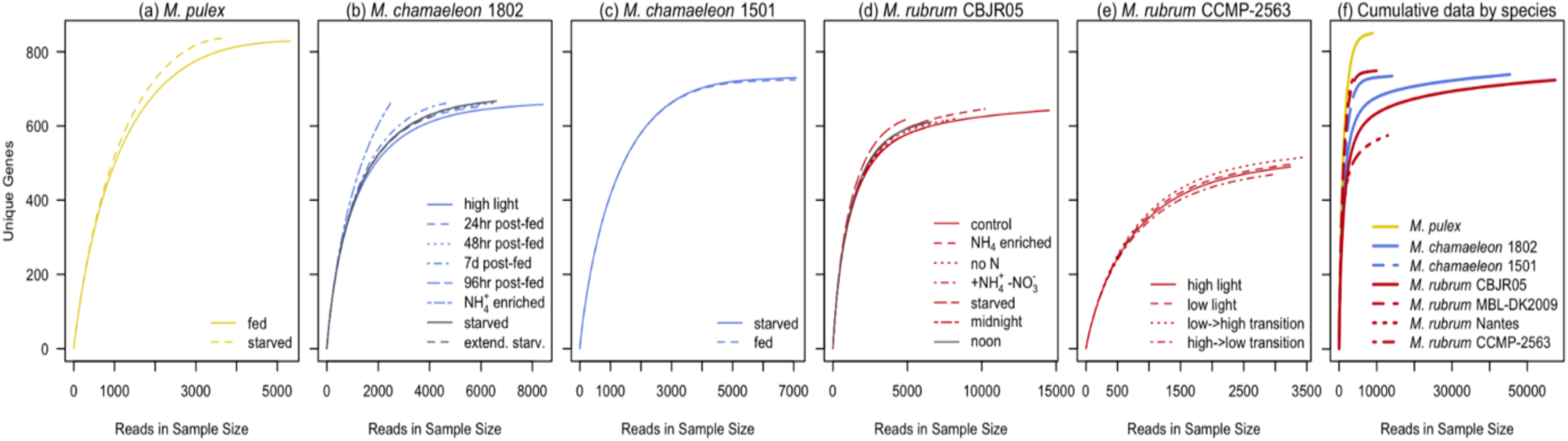
Rarefaction curves for transcriptomic data used in this study. Panels a-e show rarefaction curves by experimental treatment for transcriptomic data either generated for this study (*M. pulex*, *M. chamaeleon* NRMC-1802, and *M. rubrum* CBJR05) or previously generated by coauthors of this study (*M. chamaeleon* NRMC-1501 and *M. rubrum* CCMP-2563). Panel f shows cumulative data by species for all 7 transcriptomes used (including *M. rubrum* MBL-DK2009 and *M. rubrum* Nantes datasets generated by other labs). While all *Mesodinium* libraries appeared to saturate, *M. pulex* gene discovery saturates at a higher number of unique genes and lower number of reads than other *Mesodinium* species.

**Fig. S2.**
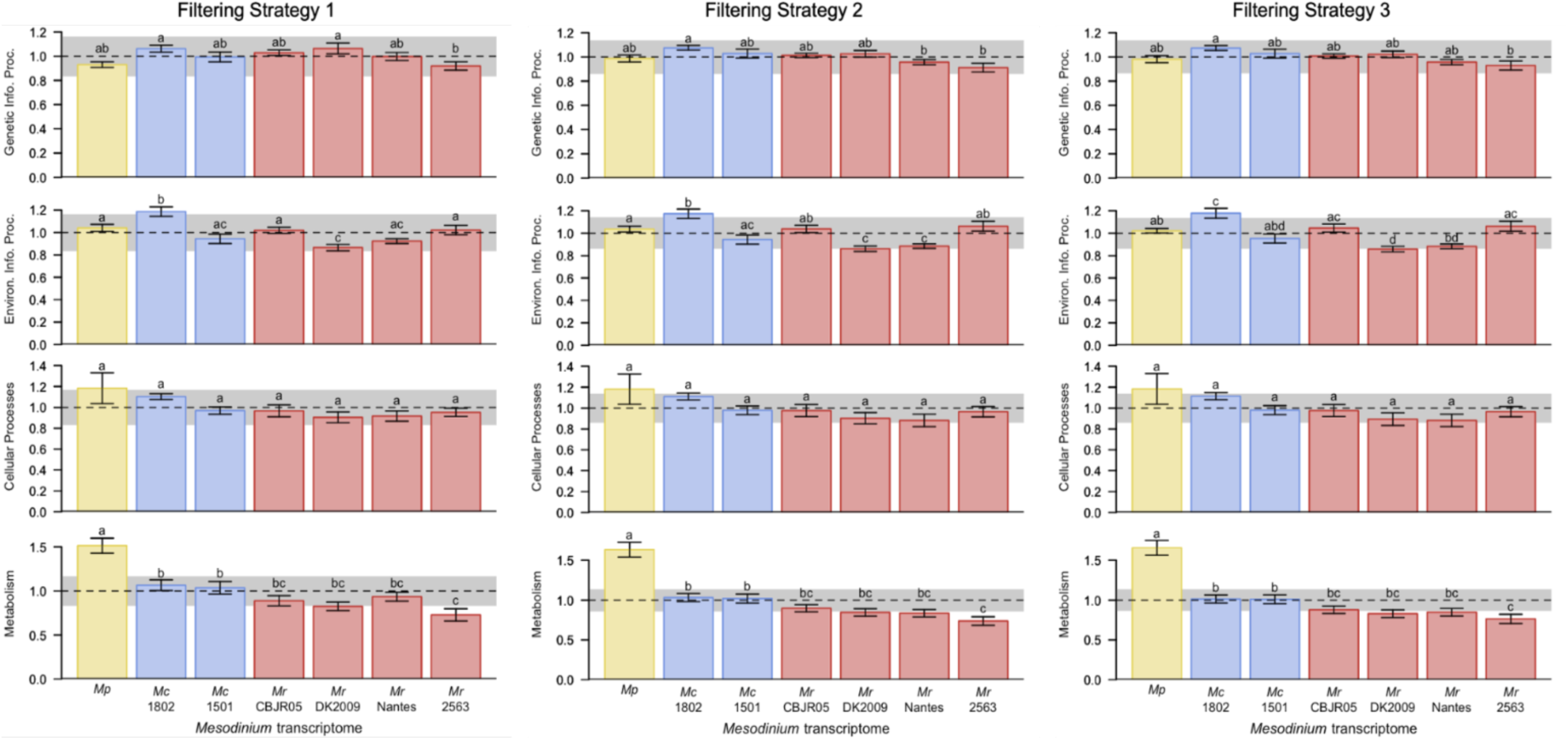
Barplots showing relative completeness for each *Mesodinium* species across different KEGG categories. For each KEGG pathway within a category, we scaled by the maximum number of unique proteins found in any *Mesodinium* transcriptome. Bar heights represent means, and whiskers represent standard deviations, of all pathways within a KEGG category. The horizontal dashed line and gray band show the mean and standard deviation of all *Mesodinium* lineages for genetic information processing, a key reference category used in our analysis. Results for three approaches to transcriptome filtering are shown.

**Fig. S3.**
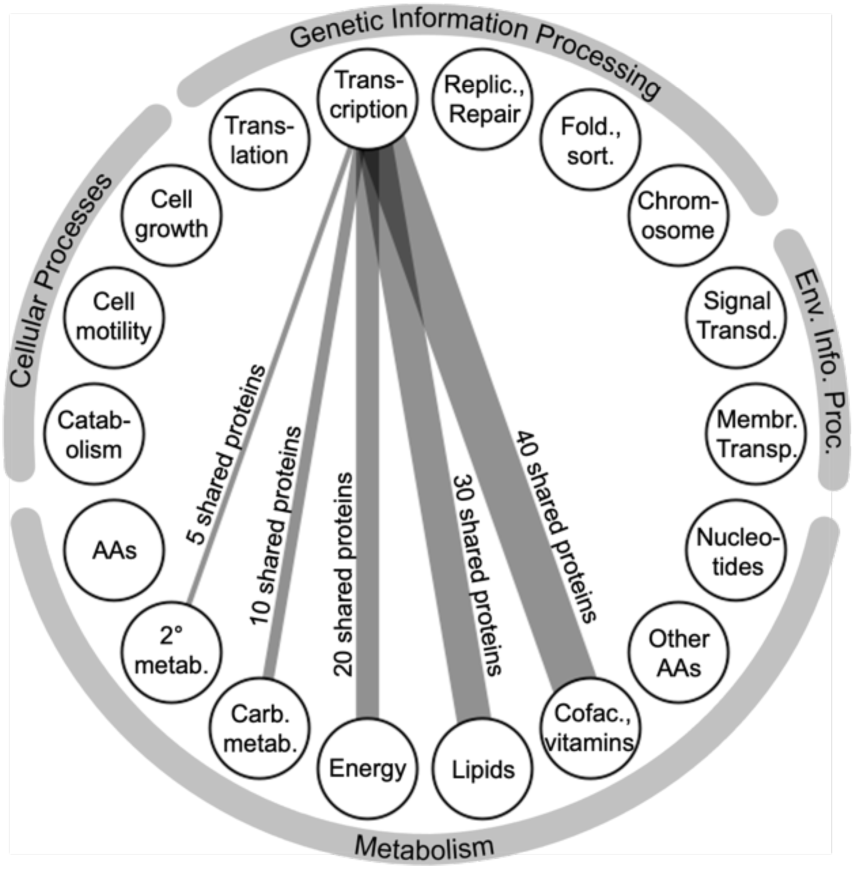
Legend for network diagrams showing KEGG subcategory node labels. A key showing weighting of connectivity by number of proteins is also included for illustrative purposes only (i.e., this does not represent actual connectivity for any lineage between Transcription and other subcategories).

**Fig. S4.**
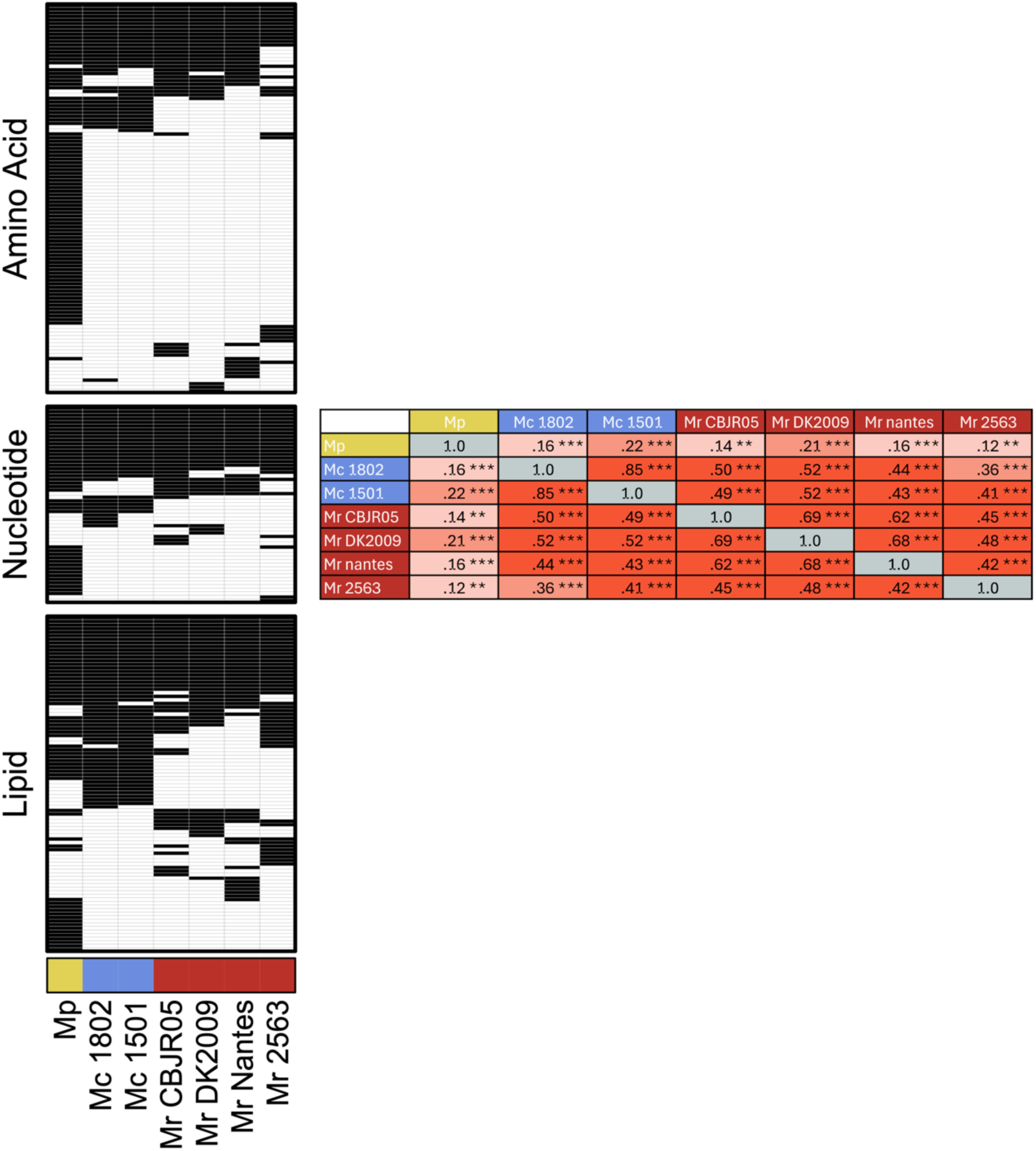
Consistency of metabolic proteins lost in *M. chameleon and M. rubrum*. a) Heatmap of proteins (KOs, rows) involved in metabolic pathways clustered by presence/absence among *Mesodinium* spp. (columns) and organized by select KEGG subcategories. b) Pairwise correlation matrix reflecting the strength of protein presence/absence patterns between *Mesodinium* spp. *** p-values << 0.01. Mp, *Mesodinium pulex* EPMP; Mc 1802, *Mesodinium chamaeleon* NRMC1802; Mc 1501, *Mesodinium chamaeleon* NRMC1501; Mr CBJRO5, *Mesodinium rubrum* CBJR05; Mr DK2009, *Mesodinium rubrum* DK2009; Mr nantes, *Mesodinium rubrum* nantes; Mr 2563, *Mesodinium rubrum* CCMP 2563.

**Fig. S5.**
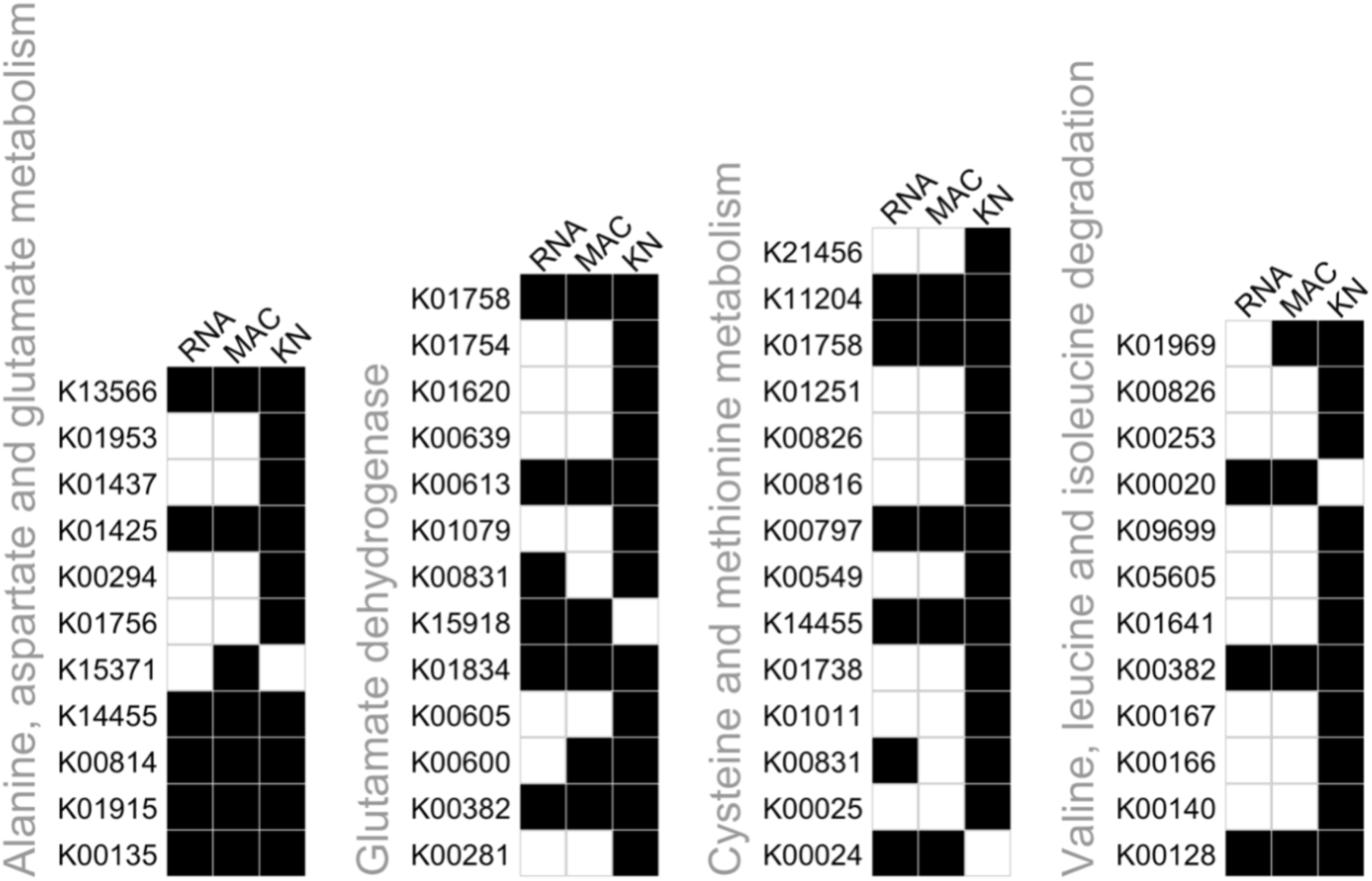
Stolen cryptophyte nuclei (“kleptokarya”) provide complementary metabolic functions associated with four major amino acid pathways. Filled boxes indicate presence, and open boxes indicate absence in the *M. rubrum* CBJR05 transcriptome (left column, “RNA”), *M. rubrum* CBJR05 macronuclear genome (middle column, “MAC”), and *T. amphioxeia*-origin kleptokaryon (right column, “KN”).

**Fig. S6.**
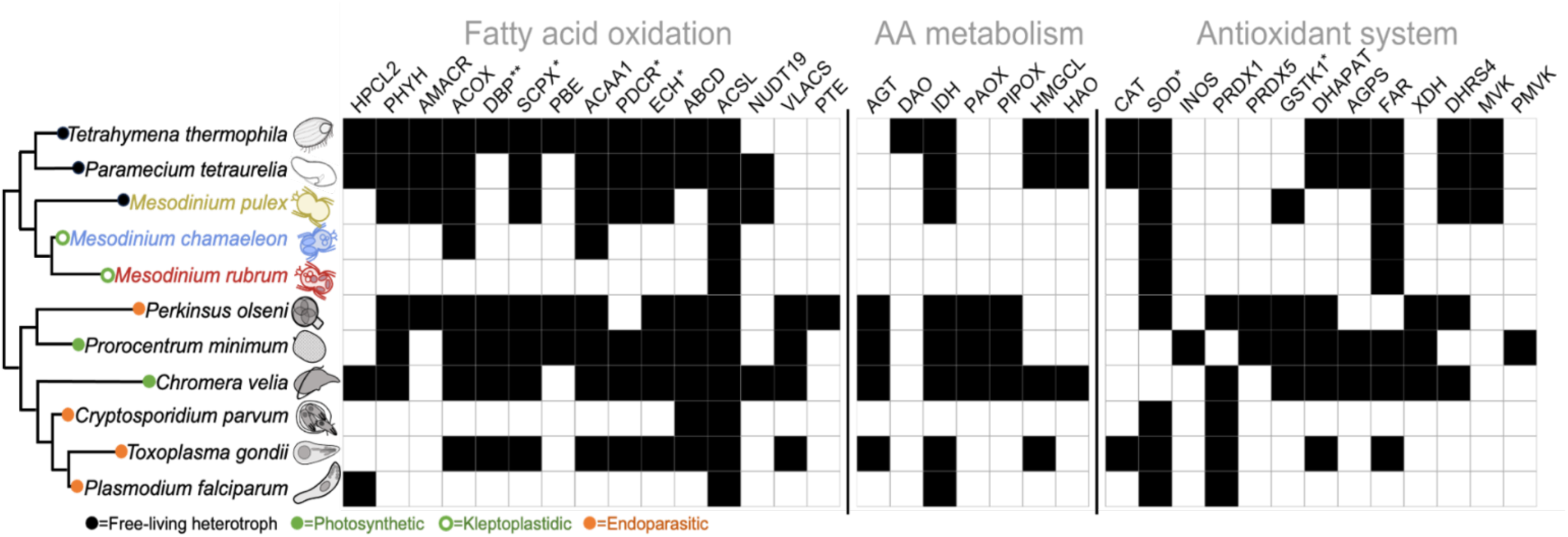
Comparison of peroxisome-related proteins in *Mesodinium* ciliates and other alveolates. Filled boxes indicate that a protein is present in a species. Data are from this study or Ludewig-Klingner et al. (*32*). The metabolic lifestyle of each lineage is indicated with a colored circle, and evolutionary relationships are shown using an unrooted dendrogram. *M. chamaeleon* data are for culture NRMC-1802, and *M. rubrum* data are for strain CBJR05. * = PTS1 or PTS2 found in *M. pulex*. ** = PTS1 or PTS2 found in *M. pulex* and *M. chamaeleon*.

**Fig. S7.**
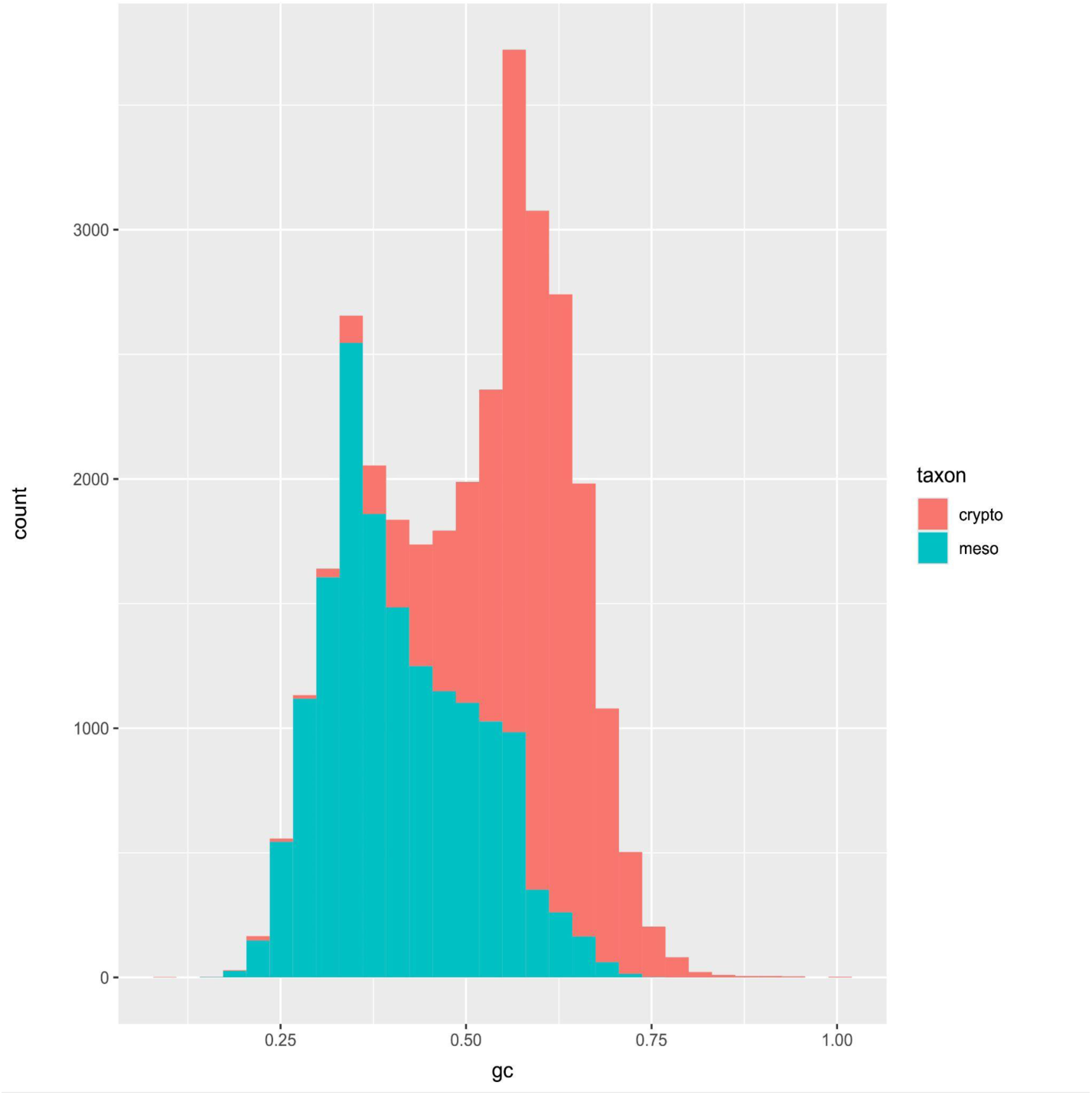
GC distribution of *Mesodinium* spp. and cryptophyte prey species coding sequences. Crypto, cryptophyte; meso, *Mesodinium* spp.

**Table S1.**
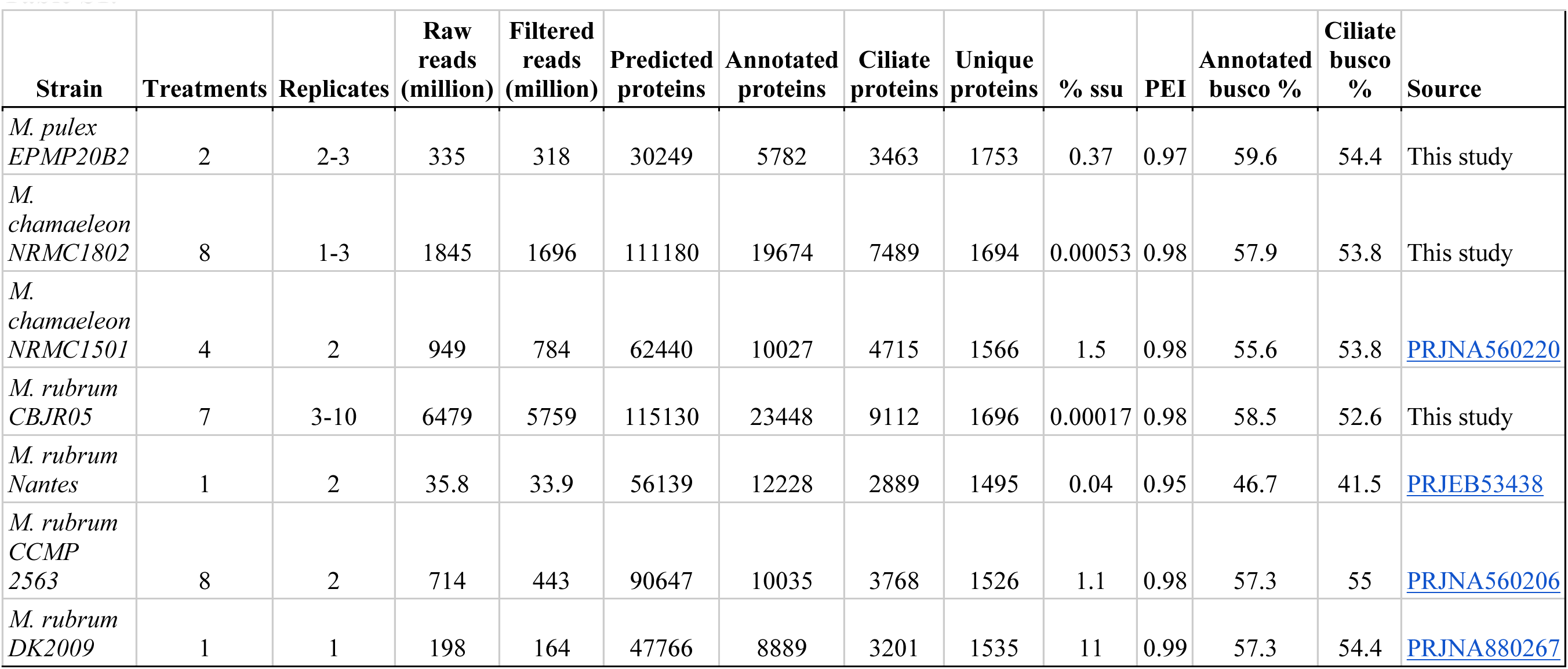
Transcriptome summary statistics. Transcriptome summary statistics. Summary statistics for seven Mesodinium RNASeq experiments and resulting transcriptome assemblies. Filtered reads, number of reads remaining after removal of bacteria and cryptophyte contamination by Kraken and BBmap, respectively; Predicted proteins, number of predicted proteins by Transdecoder after clustering at 95% identity by CD-HIT; Annotated proteins, number of proteins assigned KO numbers by GhostKOALA from KEGG; Ciliate proteins, number of annotated ciliate proteins after removal of additional bacterial and cryptophyte contamination at the protein level as well as removal of proteins with poor homology to proteins in the reference database; Unique proteins, the number of non-redundant ciliate KOs per library (representing KOs assigned to both KEGG pathways and Brite hierarchies); Annotated Busco, percent completeness of annotated transcriptome prior to removing contamination at the protein level; Ciliate Busco, percent completeness of transcriptome after removal of contamination at the protein level. Busco scores were calculated with the alveolata_odb10 database. PEI, Pielou’s Evenness Index was calculated using the read counts of the top 50 most expressed transcripts from each RNASeq library to assess the distribution of reads within *Mesodinium* spp. transcriptomes. Indices closer to 1 indicate a more even read count distribution across transcripts.

| Strain | Treatments | Replicates | Raw reads (million) | Filtered reads (million) | Predicted proteins | Annotated proteins | Ciliate proteins | Unique proteins | % ssu | PEI | Annotated busco % | Ciliate busco % | Source |
| --- | --- | --- | --- | --- | --- | --- | --- | --- | --- | --- | --- | --- | --- |
| *M. pulex*<br>*EPMP20B2* | 2 | 2-3 | 335 | 318 | 30249 | 5782 | 3463 | 1753 | 0.37 | 0.97 | 59.6 | 54.4 | This study |
| *M. chamaeleon*<br>*NRMC1802* | 8 | 1-3 | 1845 | 1696 | 111180 | 19674 | 7489 | 1694 | 0.00053 | 0.98 | 57.9 | 53.8 | This study |
| *M. chamaeleon*<br>*NRMC1501* | 4 | 2 | 949 | 784 | 62440 | 10027 | 4715 | 1566 | 1.5 | 0.98 | 55.6 | 53.8 | PRJNA560220 |
| *M. rubrum*<br>*CBJR05* | 7 | 3-10 | 6479 | 5759 | 115130 | 23448 | 9112 | 1696 | 0.00017 | 0.98 | 58.5 | 52.6 | This study |
| *M. rubrum*<br>*Nantes* | 1 | 2 | 35.8 | 33.9 | 56139 | 12228 | 2889 | 1495 | 0.04 | 0.95 | 46.7 | 41.5 | PRJEB53438 |
| *M. rubrum*<br>*CCMP 2563* | 8 | 2 | 714 | 443 | 90647 | 10035 | 3768 | 1526 | 1.1 | 0.98 | 57.3 | 55 | PRJNA560206 |
| *M. rubrum*<br>*DK2009* | 1 | 1 | 198 | 164 | 47766 | 8889 | 3201 | 1535 | 11 | 0.99 | 57.3 | 54.4 | PRJNA880267 |

## References and Notes

1. M.-A. Selosse, M. Charpin, F. Not, Mixotrophy everywhere on land and in water: the grand écart hypothesis. Ecol. Lett. 20, 246–263 (2017).

2. A. E. Douglas, Symbiosis as a General Principle in Eukaryotic Evolution. Cold Spring Harb. Perspect. Biol. 6, a016113 (2014).

3. A. L. Brown, G. A. Casarez, H. V. Moeller, Acquired Phototrophy as an Evolutionary Path to Mixotrophy. Am. Nat. 202, 458–470 (2023).

4. E. C. M. Nowack, M. Melkonian, Endosymbiotic associations within protists. Philos. Trans. R. Soc. B Biol. Sci. 365, 699–712 (2010).

5. D. K. Stoecker, M. D. Johnson, C. de Vargas, F. Not, Acquired phototrophy in aquatic protists. Aquat. Microb. Ecol. 57, 279–310 (2009).

6. S. J. Sibbald, J. M. Archibald, Genomic Insights into Plastid Evolution. Genome Biol. Evol. 12, 978–990 (2020).

7. A. Karnkowska, N. Yubuki, M. Maruyama, A. Yamaguchi, Y. Kashiyama, T. Suzaki, P. J. Keeling, V. Hampl, B. S. Leander, Euglenozoan kleptoplasty illuminates the early evolution of photoendosymbiosis. Proc. Natl. Acad. Sci. 120, e2220100120 (2023).

8. M. D. Johnson, The acquisition of phototrophy: adaptive strategies of hosting endosymbionts and organelles. Photosynth. Res. 107, 117–132 (2011).

9. J. H. Wisecaver, J. D. Hackett, Transcriptome analysis reveals nuclear-encoded proteins for the maintenance of temporary plastids in the dinoflagellate *Dinophysis acuminata*. BMC Genomics 11, 366 (2010).

10. F. Burki, B. Imanian, E. Hehenberger, Y. Hirakawa, S. Maruyama, P. J. Keeling, Endosymbiotic Gene Transfer in Tertiary Plastid-Containing Dinoflagellates. Eukaryot. Cell 13, 246–255 (2014).

11. R. G. Dorrell, C. J. Howe, Integration of plastids with their hosts: Lessons learned from dinoflagellates. Proc. Natl. Acad. Sci. 112, 10247–10254 (2015).

12. M. D. Johnson, D. Oldach, C. F. Delwiche, D. K. Stoecker, Retention of transcriptionally active cryptophyte nuclei by the ciliate *Myrionecta rubra*. Nature 445, 426–428 (2007).

13. R. Onuma, T. Horiguchi, Kleptochloroplast Enlargement, Karyoklepty and the Distribution of the Cryptomonad Nucleus in *Nusuttodinium* (= *Gymnodinium*) *aeruginosum* (Dinophyceae). Protist 166, 177–195 (2015).

14. N. Yamada, J. J. Bolton, R. Trobajo, D. G. Mann, P. Dąbek, A. Witkowski, R. Onuma, T. Horiguchi, P. G. Kroth, Discovery of a kleptoplastic ‘dinotom’ dinoflagellate and the unique nuclear dynamics of converting kleptoplastids to permanent plastids. Sci. Rep. 9, 10474 (2019).

15. A. Altenburger, H. Cai, Q. Li, K. Drumm, M. Kim, Y. Zhu, L. Garcia-Cuetos, X. Zhan, P. J. Hansen, U. John, S. Li, N. Lundholm, Limits to the cellular control of sequestered cryptophyte prey in the marine ciliate *Mesodinium rubrum*. ISME J. 15, 1056–1072 (2021).

16. M. D. Johnson, H. V. Moeller, C. Paight, R. M. Kellogg, M. R. McIlvin, M. A. Saito, E. Lasek-Nesselquist, Functional control and metabolic integration of stolen organelles in a photosynthetic ciliate. Curr. Biol. 33, 973–980.e5 (2023).

17. E. Hehenberger, R. J. Gast, P. J. Keeling, A kleptoplastidic dinoflagellate and the tipping point between transient and fully integrated plastid endosymbiosis. Proc. Natl. Acad. Sci. 116, 17934–17942 (2019).

18. M. E. S. Sørensen, V. V. Zlatogursky, I. Onuţ-Brännström, A. Walraven, R. A. Foster, F. Burki, A novel kleptoplastidic symbiosis revealed in the marine centrohelid *Meringosphaera* with evidence of genetic integration. Curr. Biol. 33, 3571–3584.e6 (2023).

19. L. F. Santoferrara, S. Guida, H. Zhang, G. B. McManus, De Novo Transcriptomes of a Mixotrophic and a Heterotrophic Ciliate from Marine Plankton. PLOS ONE 9, e101418 (2014).

20. P. J. Hansen, L. T. Nielsen, M. Johnson, T. Berge, K. J. Flynn, Acquired phototrophy in *Mesodinium* and *Dinophysis* – A review of cellular organization, prey selectivity, nutrient uptake and bioenergetics. Harmful Algae 28, 126–139 (2013).

21. H. V. Moeller, M. D. Johnson, Preferential Plastid Retention by the Acquired Phototroph *Mesodinium chamaeleon*. J. Eukaryot. Microbiol. 65, 148–158 (2018).

22. H. Dierssen, G. B. McManus, A. Chlus, D. Qiu, B.-C. Gao, S. Lin, Space station image captures a red tide ciliate bloom at high spectral and spatial resolution. Proc. Natl. Acad. Sci. 112, 14783–14787 (2015).

23. L. Guzmán, R. Varela, F. Muller-Karger, L. Lorenzoni, Bio-optical characteristics of a red tide induced by *Mesodinium rubrum* in the Cariaco Basin, Venezuela. J. Mar. Syst. 160, 17– 25 (2016).

24. A. L. Oliveira, N. Rudorff, M. Kampel, S. Sathyendranath, M. Pompeu, A. M. S. Detoni, G. M. Cesar, Phytoplankton assemblages and optical properties in a coastal region of the South Brazil Bight. Cont. Shelf Res. 227, 104509 (2021).

25. J. I. Kim, C. E. Moore, J. M. Archibald, D. Bhattacharya, G. Yi, H. S. Yoon, W. Shin, Evolutionary Dynamics of Cryptophyte Plastid Genomes. Genome Biol. Evol. 9, 1859–1872 (2017).

26. E. Lasek-Nesselquist, J. H. Wisecaver, J. D. Hackett, M. D. Johnson, Insights into transcriptional changes that accompany organelle sequestration from the stolen nucleus of *Mesodinium rubrum*. BMC Genomics 16, 805 (2015).

27. R. Albalat, C. Cañestro, Evolution by gene loss. Nat. Rev. Genet. 17, 379–391 (2016).

28. R. Poulin, H. S. Randhawa, Evolution of parasitism along convergent lines: from ecology to genomics. Parasitology 142, S6–S15 (2015).

29. M. Blume, F. Seeber, Metabolic interactions between *Toxoplasma gondii* and its host. F1000Research 7:1719 [Preprint] (2018). 10.12688/f1000research.16021.1.

30. P. J. Hansen, M. Moldrup, W. Tarangkoon, L. Garcia-Cuetos, Ø. Moestrup, Direct evidence for symbiont sequestration in the marine red tide ciliate Mesodinium rubrum. Aquat. Microb. Ecol. 66, 63–75 (2012).

31. E. Peltomaa, M. D. Johnson, *Mesodinium rubrum* exhibits genus-level but not species-level cryptophyte prey selection. Aquat. Microb. Ecol. 78, 147–159 (2017).

32. A.-K. Ludewig-Klingner, V. Michael, M. Jarek, H. Brinkmann, J. Petersen, Distribution and Evolution of Peroxisomes in Alveolates (Apicomplexa, Dinoflagellates, Ciliates). Genome Biol. Evol. 10, 1–13 (2018).

33. D. Moog, J. M. Przyborski, U. G. Maier, Genomic and Proteomic Evidence for the Presence of a Peroxisome in the Apicomplexan Parasite *Toxoplasma gondii* and Other Coccidia. Genome Biol. Evol. 9, 3108–3121 (2017).

34. T. Gabaldón, M. L. Ginger, P. A. M. Michels, Peroxisomes in parasitic protists. Mol. Biochem. Parasitol. 209, 35–45 (2016).

35. J. D. Crapo, T. Oury, C. Rabouille, J. W. Slot, L. Y. Chang, Copper,zinc superoxide dismutase is primarily a cytosolic protein in human cells. Proc. Natl. Acad. Sci. 89, 10405– 10409 (1992).

36. M. Islinger, K. W. Li, J. Seitz, A. Völkl, G. H. Lüers, Hitchhiking of Cu/Zn Superoxide Dismutase to Peroxisomes – Evidence for a Natural Piggyback Import Mechanism in Mammals. Traffic 10, 1711–1721 (2009).

37. A. Spinazzola, C. Viscomi, E. Fernandez-Vizarra, F. Carrara, P. D’Adamo, S. Calvo, R. M. Marsano, C. Donnini, H. Weiher, P. Strisciuglio, R. Parini, E. Sarzi, A. Chan, S. DiMauro, A. Rötig, P. Gasparini, I. Ferrero, V. K. Mootha, V. Tiranti, M. Zeviani, MPV17 encodes an inner mitochondrial membrane protein and is mutated in infantile hepatic mitochondrial DNA depletion. Nat. Genet. 38, 570–575 (2006).

38. J. Vasilev, A.-K. Mix, T. Heimerl, U. G. Maier, D. Moog, Inferred Subcellular Localization of Peroxisomal Matrix Proteins of Guillardia theta Suggests an Important Role of Peroxisomes in Cryptophytes. Front. Plant Sci. 13 (2022).

39. T. Cavalier-Smith, Economy, Speed and Size Matter: Evolutionary Forces Driving Nuclear Genome Miniaturization and Expansion. Ann. Bot. 95, 147–175 (2005).

40. M. Müer, Review Article: The hydrogenosome. Microbiology 139, 2879–2889 (1993).

41. N. Corradi, C. H. Slamovits, The intriguing nature of microsporidian genomes. Brief. Funct. Genomics 10, 115–124 (2011).

42. B. Striepen, A. J. P. Pruijssers, J. Huang, C. Li, M.-J. Gubbels, N. N. Umejiego, L. Hedstrom, J. C. Kissinger, Gene transfer in the evolution of parasite nucleotide biosynthesis. Proc. Natl. Acad. Sci. 101, 3154–3159 (2004).

43. P. J. Keeling, N. Corradi, H. G. Morrison, K. L. Haag, D. Ebert, L. M. Weiss, D. E. Akiyoshi, S. Tzipori, The Reduced Genome of the Parasitic Microsporidian *Enterocytozoon bieneusi* Lacks Genes for Core Carbon Metabolism. Genome Biol. Evol. 2, 304–309 (2010).

44. L. Herfort, T. D. Peterson, V. Campbell, S. Futrell, P. Zuber, *Myrionecta rubra* (*Mesodinium rubrum*) bloom initiation in the Columbia River estuary. Estuar. Coast. Shelf Sci. 95, 440– 446 (2011).

45. J. E. B. Rines, M. N. McFarland, P. L. Donaghay, J. M. Sullivan, Thin layers and species-specific characterization of the phytoplankton community in Monterey Bay, California, USA. Cont. Shelf Res. 30, 66–80 (2010).

46. K. D. Bidle, A. Vardi, A chemical arms race at sea mediates algal host–virus interactions. Curr. Opin. Microbiol. 14, 449–457 (2011).

47. L. Råberg, E. Alacid, E. Garces, R. Figueroa, The potential for arms race and Red Queen coevolution in a protist host–parasite system. Ecol. Evol. 4, 4775–4785 (2014).

48. J. J. Morris, R. E. Lenski, E. R. Zinser, The Black Queen Hypothesis: Evolution of Dependencies through Adaptive Gene Loss. mBio 3, 10.1128/mbio.00036-12 (2012).

49. A. A. Mushegian, D. Ebert, Rethinking “mutualism” in diverse host-symbiont communities. BioEssays 38, 100–108 (2016).

50. M. A. Brockhurst, D. D. Cameron, A. P. Beckerman, Fitness trade-offs and the origins of endosymbiosis. PLOS Biol. 22, e3002580 (2024).

51. E. Peltomaa, M. D. Johnson, *Mesodinium rubrum* exhibits genus-level but not species-level cryptophyte prey selection. Aquat. Microb. Ecol. 78, 147–159 (2017).

52. C. Paight, M. D. Johnson, E. Lasek-Nesselquist, H. V. Moeller, Cascading effects of prey identity on gene expression in a kleptoplastidic ciliate. J. Eukaryot. Microbiol. 70, e12940 (2023).

53. R. R. L. Guillard, J. H. Ryther, Studies of marine planktonic diatoms: I. *Cyclotella nana* Hustedt, and *Detonula confervacea* (Cleve) Gran. Can. J. Microbiol. 8, 229–239 (1962).

54. G. Cathala, J.-F. Savouret, B. Mendez, B. L. West, M. Karin, J. A. Martial, J. D. Baxter, A Method for Isolation of Intact, Translationally Active Ribonucleic Acid. DNA 2, 329–335 (1983).

55. D. E. Wood, J. Lu, B. Langmead, Improved metagenomic analysis with Kraken 2. Genome Biol. 20, 257 (2019).

56. D. Li, C.-M. Liu, R. Luo, K. Sadakane, T.-W. Lam, MEGAHIT: an ultra-fast single-node solution for large and complex metagenomics assembly via succinct de Bruijn graph. Bioinformatics 31, 1674–1676 (2015).

57. W. Li, A. Godzik, Cd-hit: a fast program for clustering and comparing large sets of protein or nucleotide sequences. Bioinformatics 22, 1658–1659 (2006).

58. W. Li, L. Fu, B. Niu, S. Wu, J. Wooley, Ultrafast clustering algorithms for metagenomic sequence analysis. Brief. Bioinform. 13, 656–668 (2012).

59. C. Camacho, G. Coulouris, V. Avagyan, N. Ma, J. Papadopoulos, K. Bealer, T. L. Madden, BLAST+: architecture and applications. BMC Bioinformatics 10, 421 (2009).

60. M. Kanehisa, M. Furumichi, M. Tanabe, Y. Sato, K. Morishima, KEGG: new perspectives on genomes, pathways, diseases and drugs. Nucleic Acids Res. 45, D353–D361 (2017).

61. B. Buchfink, K. Reuter, H.-G. Drost, Sensitive protein alignments at tree-of-life scale using DIAMOND. Nat. Methods 18, 366–368 (2021).

62. C. Jiang, S. Gu, T. Pan, X. Wang, W. Qin, G. Wang, X. Gao, J. Zhang, K. Chen, A. Warren, J. Xiong, W. Miao, Dynamics and timing of diversification events of ciliated eukaryotes from a large phylogenomic perspective. Mol. Phylogenet. Evol. 197, 108110 (2024).

63. M. D. Johnson, T. Tengs, D. W. Oldach, C. F. Delwiche, D. K. Stoecker, Highly divergent SSU rRNA genes found in the marine ciliates Myrionecta rubra and Mesodinium pulex. J. Eukaryot. Microbiol. 52, 7S–27S (2005).

64. P. Rice, I. Longden, A. Bleasby, EMBOSS: The European Molecular Biology Open Software Suite. Trends Genet. 16, 276–277 (2000).

65. E. A. Gladyshev, M. Meselson, I. R. Arkhipova, Massive Horizontal Gene Transfer in Bdelloid Rotifers. Science 320, 1210–1213 (2008).

66. C. Rancurel, L. Legrand, E. G. J. Danchin, Alienness: Rapid Detection of Candidate Horizontal Gene Transfers across the Tree of Life. Genes 8, 248 (2017).

67. R.-M. Tian, L. Cai, W.-P. Zhang, H.-L. Cao, P.-Y. Qian, Rare Events of Intragenus and Intraspecies Horizontal Transfer of the 16S rRNA Gene. Genome Biol. Evol. 7, 2310–2320 (2015).

68. M. Manni, M. R. Berkeley, M. Seppey, F. A. Simão, E. M. Zdobnov, BUSCO Update: Novel and Streamlined Workflows along with Broader and Deeper Phylogenetic Coverage for Scoring of Eukaryotic, Prokaryotic, and Viral Genomes. Mol. Biol. Evol. 38, 4647–4654 (2021).

69. B. K. B. Seah, A. Singh, D. E. Vetter, C. Emmerich, M. Peters, V. Soltys, B. Huettel, E. C. Swart, Nuclear dualism without extensive DNA elimination in the ciliate Loxodes magnus. Proc. Natl. Acad. Sci. 121, e2400503121 (2024).

70. M. Kolmogorov, J. Yuan, Y. Lin, P. A. Pevzner, Assembly of long, error-prone reads using repeat graphs. Nat. Biotechnol. 37, 540–546 (2019).

71. R. Patro, G. Duggal, M. I. Love, R. A. Irizarry, C. Kingsford, Salmon provides fast and bias-aware quantification of transcript expression. Nat. Methods 14, 417–419 (2017).

72. J. Oksanen, G. L. Simpson, F. G. Blanchet, R. Kindt, P. Legendre, P. R. Minchin, R. B. O’Hara, P. Solymos, M. H. H. Stevens, E. Szoecs, H. Wagner, M. Barbour, M. Bedward, B. Bolker, D. Borcard, G. Carvalho, M. Chirico, M. D. Caceres, S. Durand, H. B. A. Evangelista, R. FitzJohn, M. Friendly, B. Furneaux, G. Hannigan, M. O. Hill, L. Lahti, D. McGlinn, M.-H. Ouellette, E. R. Cunha, T. Smith, A. Stier, C. J. F. T. Braak, J. Weedon, Vegan: Community Ecology Package (2024; https://CRAN.R-project.org/package=vegan).

73. G. Csárdi, T. Nepusz, The igraph software package for complex network research, version 2.1.1 (2006); https://igraph.org.

74. R. Thériault, rempsyc: Convenience functions for psychology. J. Open Source Softw. 8, 5466 (2023).

75. P. K. Kim, E. H. Hettema, Multiple Pathways for Protein Transport to Peroxisomes. J. Mol. Biol. 427, 1176–1190 (2015).

76. O. I. Petriv, L. Tang, V. I. Titorenko, R. A. Rachubinski, A New Definition for the Consensus Sequence of the Peroxisome Targeting Signal Type 2. J. Mol. Biol. 341, 119–134 (2004).

